# Genome-wide association study of gastric cancer- and duodenal ulcer-derived *Helicobacter pylori* strains reveals discriminatory amino acid differences and novel oncoprotein candidates

**DOI:** 10.1101/2021.03.15.435401

**Authors:** Vo Phuoc Tuan, Koji Yahara, Ho Dang Quy Dung, Tran Thanh Binh, Pham Huu Tung, Tran Dinh Tri, Ngo Phuong Minh Thuan, Vu Van Khien, Tran Thi Huyen Trang, Bui Hoang Phuc, Evariste Tshibangu-Kabamba, Takashi Matsumoto, Junko Akada, Rumiko Suzuki, Tadayoshi Okimoto, Masaaki Kodama, Kazunari Murakami, Hirokazu Yano, Masaki Fukuyo, Noriko Takahashi, Mototsugu Kato, Shin Nishiumi, Takeshi Azuma, Yoshitoshi Ogura, Tetsuya Hayashi, Atsushi Toyoda, Ichizo Kobayashi, Yoshio Yamaoka

**Author notes:** Corresponding authors: (KY). (IK), (YY). These authors contributed equally to this work (Vo Phuoc Tuan, Koji Yahara). This author is deceased (Takeshi Azuma).

## Abstract

Genome-wide association studies (GWASs) can reveal genetic variations associated with a phenotype in the absence of any hypothesis of candidate genes. The problem of false-positive sites linked with the responsible site might be bypassed in bacteria with a high homologous recombination rate, such as *Helicobacter pylori*, which causes gastric cancer (GC). We conducted a GWAS followed by regression-based prediction of GC and duodenal ulcer *H. pylori* strains. We identified 14 single nucleotide polymorphisms (11 amino acid changes) that, combined, allowed effective disease discrimination. They were often informative of the underlying molecular mechanisms, such as electric charge alteration at the ligand-binding pocket, alteration in subunit interaction, and mode-switching of DNA methylation. We also identified three novel virulence factors/oncoprotein candidates. These results provide both defined targets for further informatic and experimental analyses to gain insights into GC pathogenesis and a basis for identifying a set of biomarkers for application in clinical settings.

## Introduction

The faster-than-exponential decrease in the cost of DNA sequencing has brought about the era of population genomics, which refers to the comparative analysis of numerous genome sequences within a population. Among the various population genomics methods available, genome-wide association study (GWAS) has the advantage of being able to reveal genetic variations associated with a particular phenotype in the absence of any hypothesis of candidate genes. GWASs have revealed the genetic basis of various human diseases, including some with multiple genetic factors. Although GWAS in bacteria has been difficult due to the strong population structure (Falush and Bowden 2006), methodological developments in the last seven years (Falush 2016; Jaillard, et al. 2018; Lees, et al. 2018) have helped to control this effect and to systematically explore relationships between a phenotypic trait and any genetic variation, such as the presence/absence of specific genes or single nucleotide polymorphisms (SNPs). GWASs utilizing these methodological developments have identified the genetic basis of several bacterial activities, including host specificity (Sheppard, et al. 2013), survival in different environments (Yahara, et al. 2017), and antimicrobial resistance (Earle, et al. 2016a; Suzuki, et al. 2016).

Berthenet et al. (2018) conducted a GWAS on *Helicobacter pylori*, a stomach pathogen that infects more than half of the global population and causes gastric cancer (GC). *H. pylori* has a very high recombination rate, leading to the generation of short recombination-derived chunks across the genome (median, 14 bp, interquartile range, 5–39 bp) (Yahara, et al. 2013). Thus, we reasoned that this feature of *H. pylori* would make it an ideal subject for a GWAS because due to the high recombination, the responsible SNP would quickly be separated from linked SNPs that otherwise would appear as false positives (S1 Figure).

The population structure of *H. pylori* is distinctive because global strains are phylogeographically differentiated and classified into several populations (hpAfrica2, hpAfrica1, hpNEAfrica, hpEurope, hpAsia2, hpSahul, and hpEastAsia) that have notable genotypic and phenotypic differences (Linz, et al. 2007); for example, the frequency of strains carrying the *cag* pathogenicity island, the strongest risk factor for GC, is nearly 100% in East Asia compared to ∼60% in other regions (Takahashi-Kanemitsu, et al. 2020). The population structure requires GWAS to be conducted separately for each population. The GWAS by Berthenet et al. (2018) was conducted in the hpEurope population. On the other hand, hspEAsia, a subpopulation of hpEastAsia, is of special interest because it has been associated with the highest incidence of GC in East Asia (Breurec, et al. 2011). This is often explained by the presence of East Asian-type CagA protein, which has distinctive sequence differences from Western CagA (Furuta, et al. 2011). CagA is encoded in the *cag* pathogenicity island and is injected into host cells, where it interacts with a number of host proteins involved in cell signaling (Hatakeyama 2017).

Molecular epidemiological studies have suggested that a single *H. pylori* virulence factor does not sufficiently explain its clinical outcomes (Yamaoka 2010). In an attempt to explore other factors, Berthenet et al. (2018) compared genome sequences of *H. pylori* strains isolated from patients diagnosed with non-atrophic gastritis (NAG), a step toward cancer, and GC patients, and reported that genes in the *cag* pathogenicity island and encoding an outer membrane protein BabA are typically associated with GC. However, the associations among genes in the *cag* pathogenicity island were as expected. In addition, the transition from NAG to GC is not discrete, but continuous, and the study did not elucidate how amino acid changes in the GWAS hits underlie the pathophysiological alterations.

Duodenal ulcer (DU) is also caused by *H. pylori*, but is considered divergent, and even somewhat mutually exclusive, from GC (Kusters, et al. 2006). DU arises from antral-predominant gastritis associated with excessive gastric acid secretion, whereas GC develops from a background of corpus-predominant gastritis or pangastritis, leading to hypochlorhydria as progressive atrophic gastritis occurs, and followed by intestinal metaplasia (Correa, et al. 1975). Cohort studies have revealed that patients with a history of DU have a considerably reduced risk of developing GC (Hansson, et al. 1996; Uemura, et al. 2001). A meta-analysis revealed that *dupA* is associated with an increased risk of DU, but a decreased risk of GC (Lu, et al. 2005). A study of DU and GC in a Moroccan population identified some specific genotypes of the virulence genes *cagA* and *vacA*s to be strongly associated with the risk of GC or DU development (El Khadir, et al. 2017). However, another study failed to detect any association between genetic factors and phenotypic traits of DU/GC (Shiota, et al. 2010). These previous studies, focusing on specific genes, did not account for the population structure of *H. pylori* strains, which would have inevitably resulted in overestimation of the extent of association (Falush and Bowden 2006). Moreover, no genome-wide study has explored a genetic marker for the discrimination of DU and GC.

In this study, we focused on DU as a reference for comparison with GC to answer the key question: what are the (potentially multiple) underlying genotypic factors in *H. pylori* that determine the risk of GC development versus DU development in infected patients and would allow differentiating between them? Unraveling these factors would be vital in understanding the microevolutionary processes toward DU/GC pathogenesis and would aid the development of clinical and epidemiological applications. To this end, we conducted a GWAS followed by regression-based prediction of a large number of *H. pylori* strains isolated from GC and DU patients in an hspEA subpopulation, and the study revealed key discriminatory amino acid differences and novel oncoprotein candidates.

## Results

### *H. pylori* strains from GC and DU in East Asia, and their population structure

We combined genome data of 209 newly sequenced strains isolated in Vietnam and Japan with those of 383 strains representing the diverse global subpopulations (Thorell, et al. 2017) and those of 22 additional strains registered in the National Center for Biotechnology Information (NCBI) GenBank repository with information on host disease status (GC or DU) (Supplementary file 1). We used the ChromoPainter/fineSTRUCTURE pipeline (Lawson, et al. 2012) to identify 11 clusters that were found in a previous study (Thorell, et al. 2017) (S2 Figure). After removing strains that were either unassigned to hspEAsia or without information of host disease status, 240 hspEAsia strains (125 GC and 115 DU) were selected. Among them, 137 isolates were from Japan, 87 from Vietnam, eight from Singapore, five from China, one from South Korea, one from Malaysia, and one from an unknown source (Supplementary file 2). A maximum-likelihood tree constructed based on core-genome SNPs revealed that, despite the substantial population structure, no cluster was solely associated with one disease status (GC or DU) (Figure 1). The tree comprised two large clusters of mostly Japanese and Vietnamese (light green and orange in the 2^nd^ column “country” in Figure 1, respectively) strains that corresponded to two hspEAsia subpopulations in the fineSTRUCTURE results (S2 Figure), although there was no significant difference in disease status frequency between these clusters (*P* = 0.2, Chi-square test).

**Figure 1.**
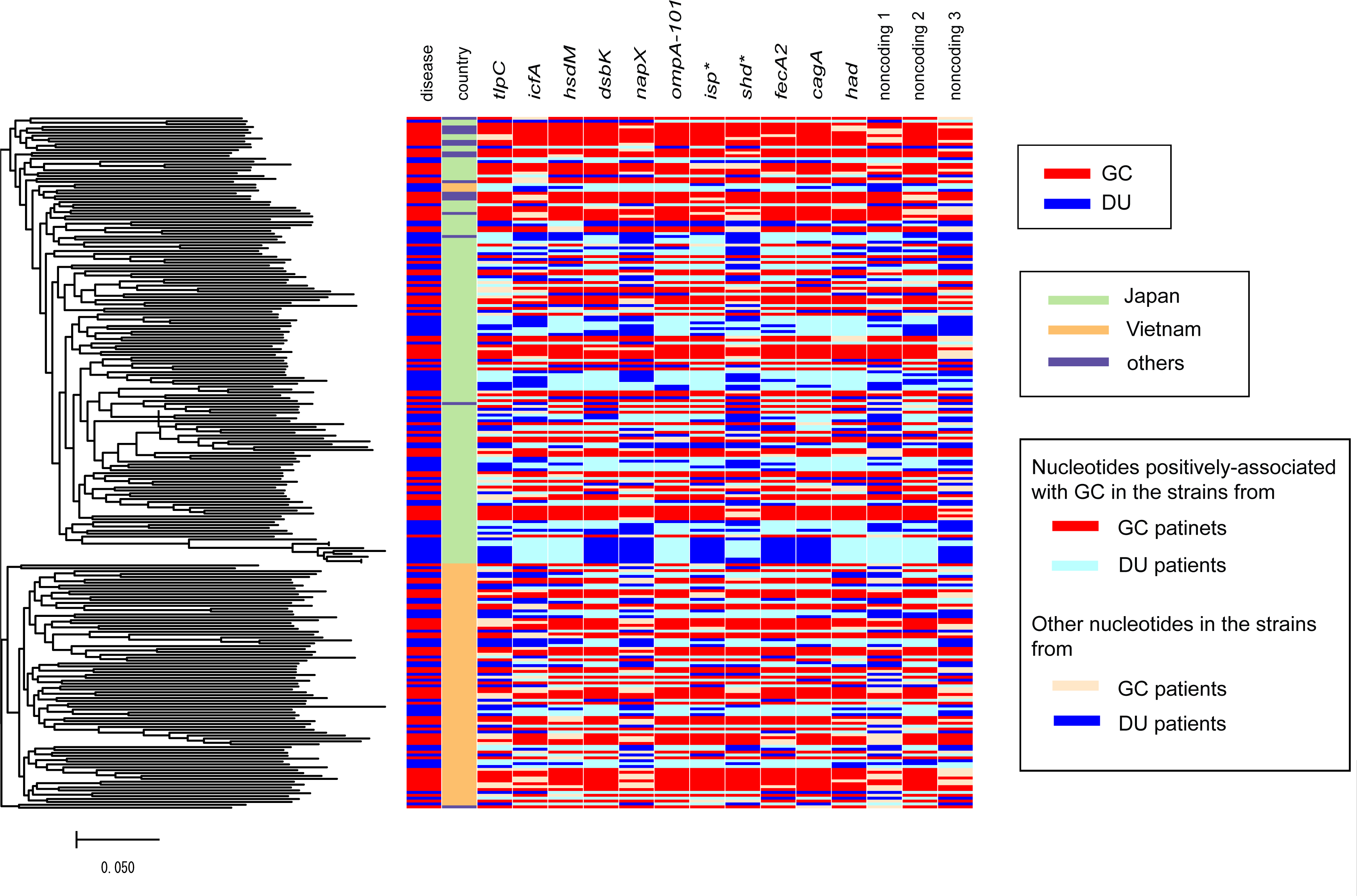
Core-genome phylogeny and metadata of the 240 strains from an hspEAsia population. Left: Mid-point rooted core-genome phylogeny. Heatmap: column 1, host disease status (DU or GC). Column 2: country of isolation. Columns 3–14 correspond to the genes carrying a nucleotide positively associated with GC.

### GWAS identifies 14 SNPs positively associated with GC

After adjustment for population structure and targeting 157,447 SNPs with a minor allele frequency >5%, the GWAS results showed that the *P*-values of most SNPs were as expected under the null hypothesis of no association (Q-Q plot in Figure 2). This indicates the absence of systematic inflation of *P*-values after adjustment for population structure (i.e., genetic relatedness among strains in hspEAsia). At the same time, we found outlier SNPs deviating from the null hypothesis (above the horizontal dashed line in Figure 2), comprising 13 non-synonymous and three intergenic SNPs (red and green dots in Figure 2) as well as 15 synonymous SNPs. Among the 16 non-synonymous and intergenic SNPs, 88% (14/16) were positively associated with GC (i.e., these SNPs had a nucleotide that showed an increase in frequency in strains isolated from GC patients compared to those isolated from DU patients), whereas the remaining 12% (2/16) had nucleotides that decreased in frequency in strains isolated from GC patients.

**Figure 2.**
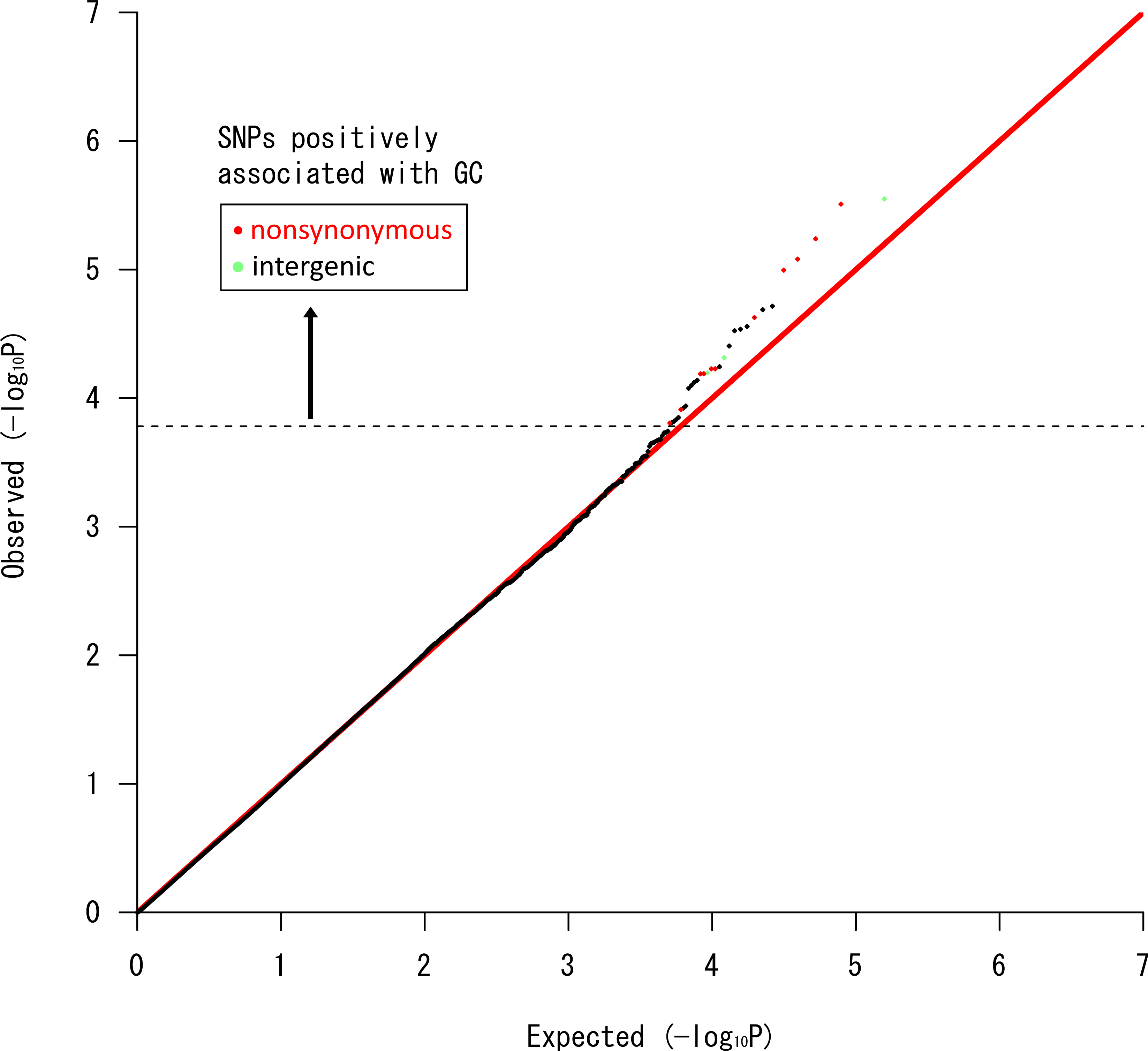
Q-Q plot to assess the GWAS results. Each dot indicates a SNP. Y-axis: observed − log_10_ (*P*) of each SNP, where *P* is its *P*-value. X-axis: expected − log_10_ (*P*) under the null hypothesis of no association. The dashed horizontal line indicates the cutoff used to judge deviation from the null hypothesis. Non-synonymous, intergenic, and synonymous SNPs satisfying the cutoff and positively associated with GC are presented as red, green, and black dots, respectively.

A simple discriminatory score calculated for each individual strain after combining information of the 14 SNPs was significantly higher among strains isolated from GC patients than among DU-derived strains (median, 38.8 versus 11.8, *P* = 2.2 × 10^−16^, Wilcoxon’s rank sum test) (Figure 3). Each SNP involved a nucleotide that showed a 12–29% frequency increase in the GC population. The phylogenetic distribution of the 14 nucleotides positively associated with GC is shown as a heatmap in Figure 1.

**Figure 3.**
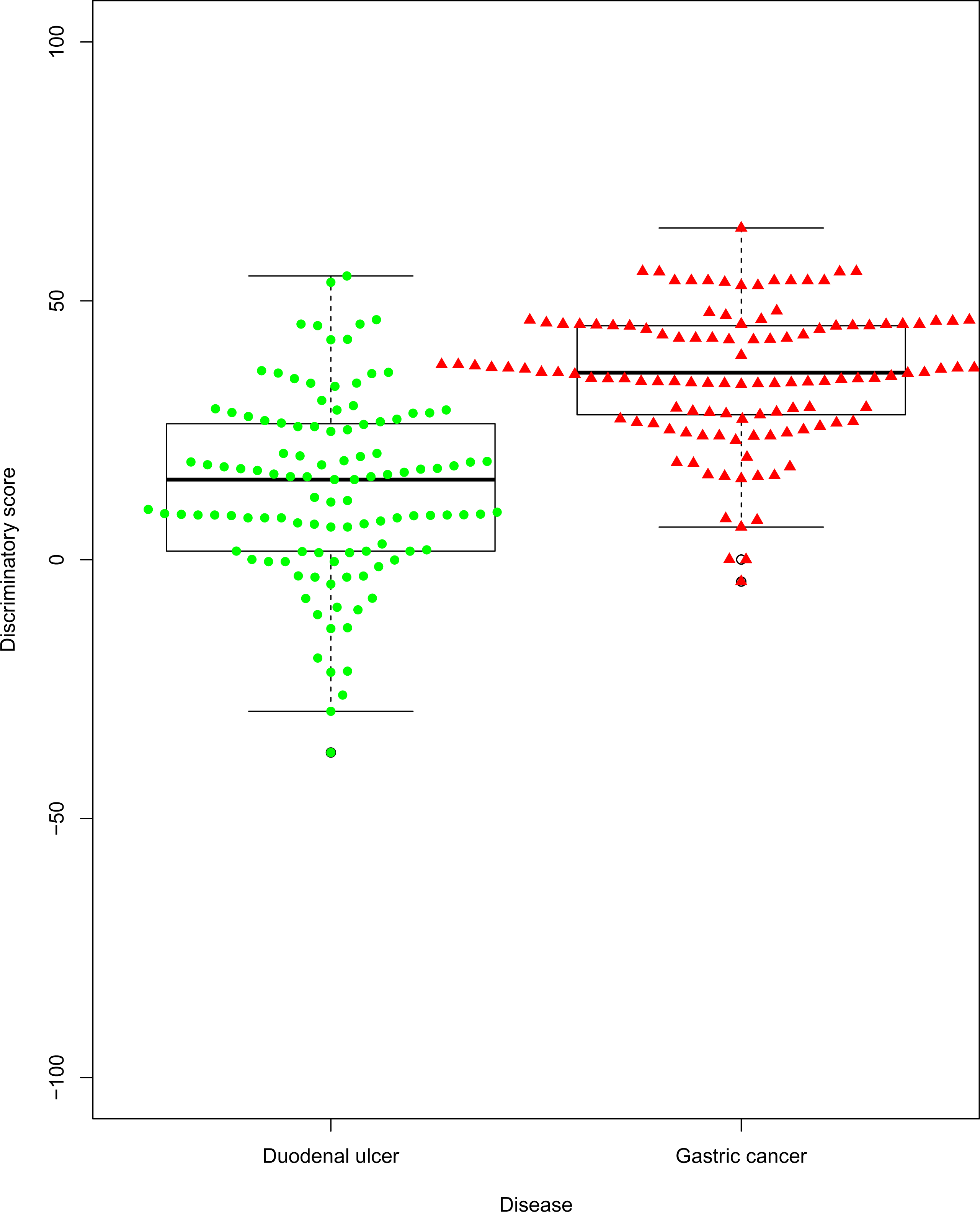
Distribution of the discriminatory score for the 240 hspEAsia strains according to host disease status. Each point corresponds to the discriminatory score of a single strain.

Eight of the 14 nucleotides positively associated with GC were derived (i.e., different from those of the European reference strain 26695 isolated from a NAG patient. Note that there is no reference genome of a European strain isolated from a DU patient). However, including only the eight nucleotides reduced the difference in discriminatory score (median, 22.34 versus 7.13). Given that the change from NAG to GC is continuous, and the presence of the remaining six nucleotides in the strain isolated from a NAG patient does not indicate prevention of GC, we did not exclude them as they are useful for discrimination between GC and DU.

A logistic regression model using the score as an explanatory variable to predict the probability of a strain being isolated from a GC patient revealed that the area under the receiver operating characteristic (ROC) curve (AUC) indicating discriminatory capacity was 0.88 (95% CI, 0.81–0.95) (S3A Figure) when we conducted a cross-validation in which the first half of strains in the maximum-likelihood tree (Figure 1) was used to train the model and the second half for prediction. We obtained similar results by two other means of cross-validation: AUC 0.86 (95% CI, 0.79–0.93) when the Japanese strains were used for training and the remaining strains for prediction (S3B Figure), and AUC 0.93 (95% CI, 0.89–0.97) when the Vietnamese strains were used for training and the remaining strains for prediction (S3C Figure). These results indicated the high discriminatory performance of the 14 SNPs combined.

In addition, we conducted a GWAS using another program, pyseer, which can adjust for population structure, yielding *P*-values that were highly correlated with those obtained above (Spearman’s correlation coefficient, 0.89). Again, most *P*-values were as expected under the null hypothesis of no association, whereas there were more outlier SNPs deviating from the null hypothesis, including all the above 14 SNPs (pink dots in the Q-Q plot in S4 Figure).

Finally, we conducted a GWAS to assess the presence or absence of a specific gene among 10,499 genes identified via pan-genome analysis using the Roary pipeline (Page, et al. 2015). However, for most genes, the *P*-values were not as expected under the null hypothesis of no association, indicating that the population structure was not well adjusted (Q-Q plot in S5 Figure). This result indicated that not genes, but SNPs are an appropriate unit for GWAS in *H. pylori*.

### Amino acid changes suggest molecular mechanisms underlying the disease phenotype

The genes harboring the 11 non-synonymous SNPs and those closest to the three intergenic SNPs are shown in Table 1 (ordered according to the y-axis in the Q-Q plot in Figure 2). We placed the amino acid changes on predicted protein 3D structures to analyze their function and biological significance. Unexpectedly, this process provided deep insights into the molecular mechanisms underlying the different diseases, as illustrated in Figures 4 and 5, and S6 Figure.

**Table 1.** SNPs positively associated with GC identified by GWAS.

| Genomic position | Locus including or closest to the SNP | Gene name | Description | Function | Nucleotide associated with GC | Nucleotide position in the gene | Amino acid change | Corresponding locus in the strain 26695 |
| --- | --- | --- | --- | --- | --- | --- | --- | --- |
| 533482 | 91 bp upstream of HPF57_0521 [532255..533391] (-) | *kch, trkA* | potassium channel | potassium conductance regulator | G | -91 |  | HP0490 |
| 256111 | HPF57_0250 [255679..256476] (+) | *dsbG/K* | thiol:disulfide interchange protein | disulfide bond formation for secretion | A | 433 | K145E | HP0231 |
| 155782 | HPF57_0151 [155438..156307] (+) | *triH\** | BIR, Dps/NapA, RAD21 similarity | host interference? | A | 345 | K115K,X | HP0130 |
| 96796 | HPF57_0094 [95423..96958] (-) | *tlpC* | chemotaxis sensor | chemotaxis | A | 163 | K55E,Q | HP0082 |
| 1459449 | HPF57_1382 [1459397..1460092] (+) | *ompA101* | outer membrane protein | transport and structure | T | 53 | V53A | HP1467 |
| 1434839 | HPF57_1355 [1434577..1435371] (-) | *isp\** | DegS/HtrA homolog | inhibitor of | G | 533 | G178E | HP1440 |
| 904207 | 14bp downstream of HPF57_0865 | *rpoZ* | DNA-directed RNA polymerase subunit omega | prophage silencing? stringent response? | A |  |  | HP0843 |
| 1296088 | HPF57_1209 [1295855..1296457] (-) | *csd5* | cell shape determinant | helical cell shape | A | 370 | N124H,Y,D | HP1250 |
| 871135 | HPF57_0829 [870769..873144] (-) | *fecA2* | iron importer | iron uptake | C | 2010 | S670X,S | HP0807 |
| 839132 | 29bp upstream of HPF57_0798 [838879..839103] (-) | *thiE* |  | supplying vitamin B1? | A |  |  | HP0776 |
| 598098 | HPF57_0574 [594999..598511] (+) | *cagA* | cytotoxin-associated gene A | interference with signal transduction | G | 3100 | A1034T,X,S | HP0547 |
| 146365 | HPF57_0139 [145564..146508] (-) | *ctbP\** | human C-terminal binding protein homolog | interference with gene expression? | G | 144 | E48EDX | HP0096 |
| 2253 | HPF57_0004 [1728..2393] (-) | *icfA* | beta-carbonic anhydrase | acid acclimation | G | 141 | L47L,X | HP0004 |
| 498631 | HPF57_0490 [497344..498975] (-) | *hsdM* | Type I restriction enzyme M protein | DNA methyltransferase | T | 345 | D115D,X | HP0463 |

position in F57 reference genome
\* designated in this work. *triH*: triple halves. *isp*: inactive serine protease. *shd*: SH3 domain. *ctbP*: C-terminal binding protein.

**Figure 4.**
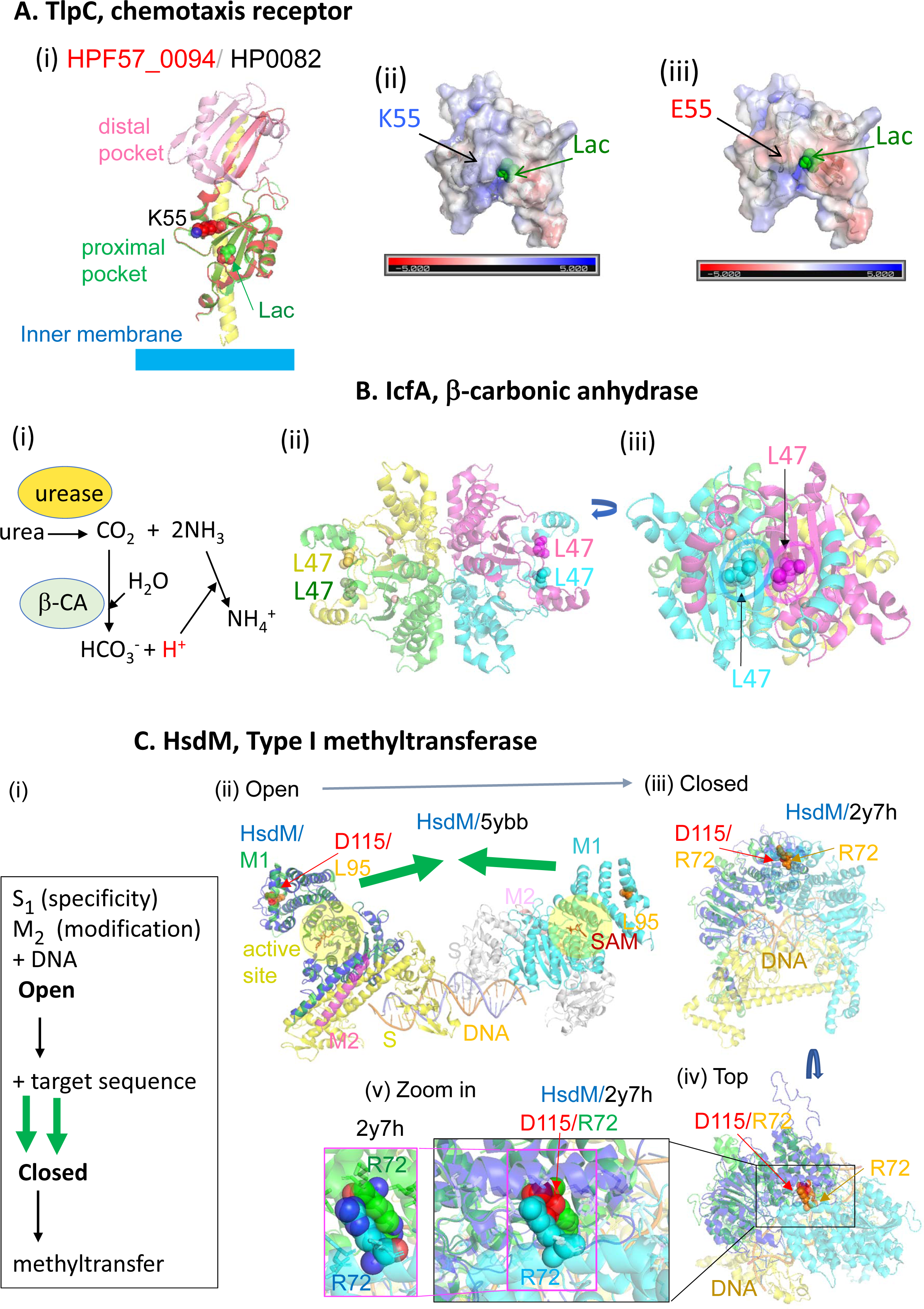
Predicted structures of proteins with discriminatory non-synonymous SNPs. (A) TlpC (HPF57_0094). (i) Model on the homolog of strain 26695 (PDB 5wbf) (Machuca, et al. 2017). K55 in HPF57_0094 corresponds to E217 in HP0082, which is split into two genes in the Japanese reference strain F57. (ii)–(iii) Surface electric charges. E55 mutant protein was generated from the model by mutagenesis (PyMOL). (B) IcfA (HPF57_0004). (i) Acid acclimation in the cytoplasm together with urease. (ii)–(iii) HPF57_0004 on *E. coli* homolog (PDB 1i6p). (C) HsdM (HPF57_0490). (i) Reaction steps of a Type I restriction-modification enzyme (Liu, et al. 2017). (ii) Model on 5ybb in PDB, two methyltransferase molecules, each 2M + 1S, of *Caldanaerobacter subterraneus*. D115 corresponds to L95. (iii)–(v) Model on 2y7h in PDB, a model of EcoKI methyltransferase based on EMD-1534 (Kennaway, et al. 2009). D115 corresponds to R72.

**Figure 5.**
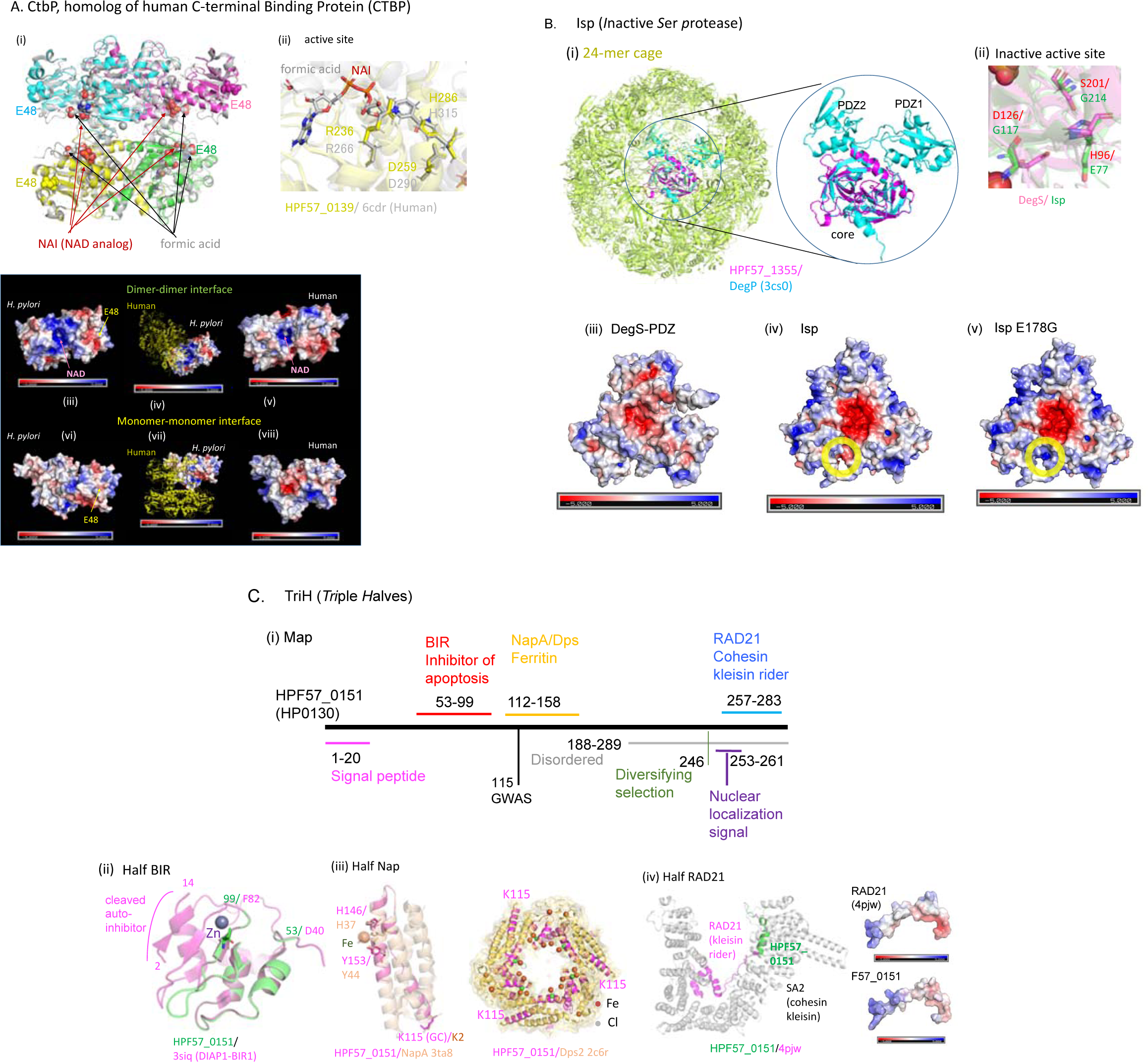
Predicted structures of four new virulence factor/oncoprotein candidates. (A) CtbP, C-terminal binding protein (HPF57_0139, HP0096). (i) Model on human CtBP1 (PDB 6cdr). (ii) Similarity of their active sites. (iii) Subunit interaction and surface electric charges in a model on 3kb6 (*Aquifex aeolicus* homolog) and CtBP1. (B) Isp (*i*nactive *S*er *p*rotease). (i) F57_1355 modeled on *E. coli* DegP. (ii) Active site of F57_1355 modeled on 3lgi.1 in PDB (*E. coli* DegS). The three amino acids (HDS triad) responsible for activity are all replaced. (iii)–(v) Surface electric charge distribution in *E. coli* DegS without PDZ (Sohn, et al. 2010) (3lgi.1 in PDB), HPF57_1355 modeled on it, and the E178G mutant generated *in silico*. (C) TriH, Triple halves. HPF57_0151. HP0130. (i) Map. “Disordered” is from UniProt. Nuclear localization signal is by cNLS Mapper. “Diversifying selection” is from a previous study. (ii)–(iv) Similarity to three half domains. (iii) Modeled on NapA (strain YS39, 4evd in PDB) and aligned with iron-soaked NapA (YS39, 3ta8 in PDB). Fe-interacting residues as well as the GWAS residues are in sticks. 2c6r in PDB is Dps2 in *Deinococcus radiodurans*. Note the difference in NapA coordinates in the literature (Zanotti, et al. 2002; Tsuruta, et al. 2012). (iv) HPF57_0151 modeled on PDB 4pjw (human).

TlpC (HPF57_0094), a chemotaxis receptor (Figure 4A (i)), has an extra-cytoplasmic ligand-binding domain. TlpC binds to lactate, which is known to promote *H. pylori* growth in the stomach, as an attractant (Machuca, et al. 2017). The structure of its proximal pocket in the Japanese reference strain F57 isolated from a GU patient was predicted to be very similar to that in the solved structure of HP0082 for the European reference strain 26695, which contains a bound lactate. Amino acid substitution at position 55 (Table 1), corresponding to the discriminatory SNP, alters the electric charge distribution around the ligand-entry site (Figure 4A (ii) (iii)) and may affect its ligand-binding properties, e.g., on-rate, off-rate, or affinity. These types of changes could influence the types of responses, and may in turn influence *H. pylori* growth or survival. Lactate was recently identified as a highly used molecule in the stomach, and one that varies between stomach regions (Keilberg, et al. 2020).

The α family of carbonic anhydrases in the periplasm processes CO_2_ produced from urea by urease and is essential for acid acclimation (Supuran and Capasso 2016). The β family of carbonic anhydrases, including IcfA (HPF57_0004, Figure 4B (i)), likely plays the same role in the cytoplasm (Nishimori, et al. 2007). Residue L47, corresponding to the discriminatory SNP, forms a pair at the dimerization joint (Figure 4B (ii) (iii)). We predicted that this residue affects subunit interaction to regulate the reaction speed of this very fast-acting enzyme. It could be related to the elevated acid secretion in UCs and its suppression during GC progression.

One GWAS SNP was mapped on a DNA methyltransferase, M subunit of a Type I restriction-modification system (HPF57_0490, Figure 4C). In addition to being responsible for modification against restriction, these methyltransferases affect gene expression (Yano, et al. 2020). When the two assemblies of the methyltransferase (2 × S1M2) recognize their target DNA sequences, they transform from an open to a closed form (Liu, et al. 2017). The amino acid residue D115 on the helices of the two assemblies moves a large distance to bind and thus connect the two assemblies. A mutation in the *E. coli* homolog corresponding to the residue at position 115 and some mutations in this helix switch the enzyme from maintenance methylation mode (hemi-methylated to fully methylated DNA) to *de-novo* methylation mode (unmethylated to fully methylated DNA) (Kelleher, et al. 1991), somehow recognizing the methylation status of the target DNA sequence. This likely affects restriction attacks on incoming unmethylated DNA and on endogenous DNA upon loss of methylation. The necessity of such destructive genome maintenance is expected to differ between the stomach environment and the duodenal environment as well as between different ulcer/cancer stages. Amino acid changes corresponding to the discriminatory SNPs included four more known virulence factors: CagA oncoprotein (S6A Figure), DsbG/K disulfide bond (S-S)-forming enzyme (S6B Figure), FecA-2 iron importer (S6C Figure), and OmpA101 outer membrane protein (S6D Figure).

### Novel virulence factor/oncoprotein candidates

Our GWAS identified three new virulence factor/oncoprotein candidates. The first (HPF57_0139, HP0096), designated CtbP (*C*-*t*erminal *b*inding *p*rotein) here, is similar to the human “C-terminal of adenovirus E1A” binding protein 1 (CtBP1) (34% amino acid sequence identity in BLAST) and CtBP2 (32%) (Bellesis, et al. 2018) in sequence and in structure (GMQE = 0.69 and QMEAN = –1.99 on CtBP1 (6cdr in PDB) and GMQE = 0.66 and QMEAN = –2.42 on CtBP2 (4lcj in PDB) in SWISS-MODEL) (Figure 5 (i), (ii)). This family (NAD-dependent D-isomer specific 2-hydroxyacid dehydrogenase, CH-OH + NAD => C=O + NADH) is prevalent. Human CtBPs are NAD-sensing transcriptional co-regulators that bind to transcription factors and recruit chromatin-remodeling enzymes to gene promoters (Chinnadurai 2009). Their dehydrogenase activity with NAD as the acceptor is used to monitor intracellular NADH/NAD concentrations (or the energy state, in a sense). Both CtBPs have been implicated in repression of the epithelial phenotype and of apoptotic pathways, and in cancer progression (Chinnadurai 2009). The *H. pylori* homolog has a more basic charge around the NAD-binding site (Figure 5A (iii) vs. (v)) and a stronger dimer-dimer interaction (Bellesis, et al. 2018). *H. pylori* CtbP might interfere with NAD sensing and/or assembly of human CtBPs and affect their NAD-based transcription regulation and cell death/differentiation. *H. pylori* may inject its CtbP into the host cell to hack it at the CtBP hubs of a protein-protein network, just as adenovirus uses E1A (King, et al. 2018). V65 in CtBP1, corresponding to GWAS amino acid E48 in *H. pylori* CtbP, is located at the N-terminal region (1–70), which likely binds to GLIS2 (GLI-similar 2), a Krüppel-like zinc finger transcription factor that maintains the differentiated epithelial phenotype in renal cells (Jetten 2019).

The second new virulence factor candidate is Isp (*i*nactive *s*erine *p*rotease), HPF57_1355 (HP1440) (Figure 5B). It is structurally similar to the HtrA (*h*igh-*t*emperature *r*equirement) family of serine proteases/chaperones (i), which maintains protein quality in the periplasmic or intermembrane spaces of mitochondria in animals and plants. *H. pylori* HtrA cleaves cell-to-cell junction factors and extracellular matrix proteins, disrupting the epithelial barrier (Backert, et al. 2018). Human HtrAs modulate mitochondrial homeostasis, cell signaling, and apoptosis, and disturbances in their action are linked to oncogenesis and neurodegeneration (Zurawa-Janicka, et al. 2017).

Although Isp and HtrA family sequences are not very similar, their structural similarity is high (QMEAN = –2.1 of 39–124 in Swiss-Prot with mitochondrial serine protease HtrA2, 5TO0 in PDB). The predicted structure of Isp shows its important feature, i.e., concentration of negative charges on one side of the homo-trimer joint (funnel) (iv)–(v). However, its expected active site lacks all three amino acids (HDS triad) in the active site (ii). It also lacks the arm-like PDZ domains (Wilken, et al. 2004) (i). It carries a signal peptide (aa 1–24, UniProt) as a bacterial DegP/Q subfamily, which includes *H. pylori* HtrA. Figure 5B (i) shows Isp modeled on and aligned with a 24-mer cage of DegP (Backert, et al. 2018). The negatively charged area in the funnel is narrowed by an amino acid substitution at position 178, corresponding to the discriminatory SNP (Figure 5B (iv)–(v)), presumably affecting protein binding. We expect that this protein interferes with HtrA family proteases in human mitochondria, other bacteria, or their own (HtrA).

The third virulence factor/oncoprotein candidate is TriH (*tri*ple *h*alves) (HPF57_0151, HP0130), which carries three half pathogenicity-related domains (Figure 5C). It carries a signal peptide (1–20) (SignalP 5.0) and is likely secreted. It also carries a strong nuclear localization signal, KPKKKRRLS (cNLS Mapper although nuclear localization was not experimentally verified (Lee, et al. 2012). The first half domain (aa 53–99) is the inhibitor of apoptosis (BIR) domain (Li, et al. 2011) (QMEAN = –1.6 in SWISS-MODEL), which inhibits a caspase (Figure 5C (ii)) (Li, et al. 2011); thus, this domain may interfere with apoptosis. The second domain (112–158 aa) corresponds to half of a ferritin, belonging to the Dps (DNA-protecting proteins under starved conditions) family (Figure 5C (iii)) (QMEAN = –1.5 on PDB 2c6r), which stores Fe inside 12-mer shells and protects DNA from oxidative damage. Dps of *H. pylori*, designated NapA (neutrophil-activating protein) (Zanotti, et al. 2002), attracts neutrophils, promotes their adhesion to endothelial cells, and induces oxygen radical production. The similar region consists of two helices out of the four-helix bundles and contains only two of the four metal-binding residues (Zanotti, et al. 2002) (Figure 5C (iii)). Residue K115, corresponding to the discriminatory SNP, corresponds to K2 on NapA at the end of a helix. This region might modify NapA action.

The C-terminal part of TriH is disordered (UniProt) and has a site of diversifying selection (Figure 5B) (Yahara, et al. 2016). The amino acid region 257–283 is similar in sequence, structure, and electric charge to a part of RAD21/Scc1, cohesin kleisin rider (Figure 5C (iv)) (QMEAN = –0.15). A cohesin ring encircles and coheres replicated chromosomes until its cleavage triggers metaphase-to-anaphase transition. In addition, it generates, maintains, and regulates intra-chromosomal DNA looping events. Cohesin kleisin SA2 cleaves and seals the cohesin ring under regulation by its rider, RAD21. Cohesin is among the most commonly mutated protein complexes in cancer (Waldman 2020), and somatic mutations and amplification of RAD21 have been reported in human tumors (Cheng, et al. 2020). The N-terminus of the corresponding sequences (321–346 in RAD21) contains part of the nuclear localization signal: 317–339 in RAD21 (Cheng, et al. 2020) and 253–261 in TriH. We expect that this TriH domain enters nuclei and interacts with cohesin kleisin to affect cohesin action.

Our fourth novel virulence factor candidate, HPF57_1209 (HP1250), with an SH3b domain, turned out to be the previously studied Csd5 for helical cell shape (Blair, et al. 2018). Its C-terminal SH3b domain binds to peptidoglycan, while its N-terminal membrane domain interacts with bactofilin, a peptidoglycan precursor synthase, and the inner membrane-spanning F1FO ATP synthase. Its N-terminal 1–30 aa is predicted to be a signal peptide (SignalP 5.0). Residue 124 in its central disordered region, corresponding to the discriminatory SNP, varied among strains. Deletion of the disordered region resulted in reduced side curvature. We hypothesize that this residue and this domain may adjust cell curvature to its target mucous layers in different tissues (stomach vs. duodenum) or to different cancer progression stages, as seen with many other proteins (Blair, et al. 2018).

### Intergenic SNPs

One of the three intergenic SNPs was found 91 bp upstream of HPF57_0521 (corresponding to HP0490) (S7 Figure). It had the lowest *P*-value (0.000003) among the discriminatory SNPs (upper right green dot in the Q-Q plot in Figure 2). Upstream of HP0490, there is an extended Pribnow box (tgnTAtaAT) as the –10 motif of sigma 80 preceded by periodic AT-rich sequences, although a transcription start site was not detected in previous experiments (Bischler, et al. 2015), likely because of the high transcription of the upstream ribosome protein gene, HP0491. In a predicted secondary structure (M-fold) of the expected transcript, the discriminatory SNP is located at a loop-stem boundary, presumably slightly affecting interaction with a protein or an RNA. Use of the sub-optimal UUG start codon instead of AUG and the presence of this antisense RNA suggest tight regulation of HP0490. The HP0490 gene (*kch, trkA*) encodes a K^+^ channel protein regulating K^+^ conductance across the membrane (UniProt) and is essential for *H. pylori* colonization of the murine stomach (Stingl, et al. 2007). Mutation of HP0490 might modulate the expression of K^+^ conductance for persistence in the gastric/duodenal environment.

The second intergenic SNP is present in the promoter region (–29 bp) of the omega subunit of RNA polymerase (HPF57_0798). (In *H. pylori*, the upstream promoter element is characterized by an AT-rich sequence (Sharma, et al. 2010).) *E. coli* omega binds ppGpp alarmon (Mechold, et al. 2013) in stringent response(Hauryliuk, et al. 2015), although its N-terminal MAR motif for ppGpp-binding is not conserved in *H. pylori* (Hauryliuk, et al. 2015). *E. coli* omega affects the transcription of prophage genes not bound by the H-NS silencer (Yamamoto, et al. 2018). Prophage action may differ between DU and GC as mentioned above for restriction-modification systems.

The third intergenic SNP at 904207 in F57 lies 14 bp downstream of an operon-like gene cluster for vitamin B1 (thiamine diphosphate) synthesis, *thiM-thiD-thiE* (HPF57_0867-HPF57_0865, HP0845-HP0843). It disrupts the stem GGAAUU/CCUUAA of the first stem-loop, and might affect the expression of these genes and vitamin B1 synthesis. Thiamine derivatives bind directly to mRNA to regulate gene expression (riboswitch) in bacteria (Winkler, et al. 2002). Vitamin B1 availability may differ between the stomach and duodenum and between cancer cells and other cells. *H. pylori* may even supply vitamin B1 to human cells to affect their growth as the microbiome contributes to vitamin metabolism (Rodionov, et al. 2019).

## Discussion

This was the first GWAS to reveal GC-related genetic features by focusing on the highest-risk *H. pylori* population of GC and to utilize the largest dataset of *H. pylori* strains isolated from GC and DU patients reported to date. Generally, it becomes more difficult to isolate *H. pylori* as the cancer stage progresses. The dataset is peerless in that is covers more than 100 strains from GC patients, as well as those from DU patients, which enabled the unprecedented GWAS. However, we should keep in mind that the sample size is still smaller than that of GWASs in other bacterial species that utilized thousands of genome sequences (Chewapreecha, et al. 2014; Earle, et al. 2016b; San, et al. 2019). Accordingly, the statistical power was insufficient to judge whether each discriminatory SNP indeed has a significant effect. When we conducted a standard multiple-testing correction, the false discovery rate was at least 0.24 for the most significant SNP. Further studies, if possible, with larger sample sizes, are warranted to test the effect of each SNP.

Thus, we rather examined the combined effect of all SNPs deviating from the null hypothesis in the Q-Q plot. To this end, we used a simple score to combine the effects of the SNPs and simple univariate logistic discriminatory analysis. It was remarkable that such a simple approach worked well (with AUC > 0.85 in all three cross-validations). Although cross-validation was conducted because of the small sample size, further studies are warranted to prepare another independent dataset of *H. pylori* isolated from GC and DU patients and utilize it to statistically validate whether discrimination using the SNPs works well in another external population.

For gastric colonization by *H. pylori*, acid acclimation in the acidic environment of the stomach is key (Peek and Blaser 1997). One of the non-synonymous discriminatory SNPs was found in *icfA* (Figure 4B), encoding an enzyme involved in gastric pH homeostasis. Indeed, DU commonly develops under a low pH condition, in contrast to GC (Kusters, et al. 2006).

Another discriminatory SNP was identified in FecA-2 (S6C Figure), involved in iron uptake. Iron uptake and metabolism are central to *H. pylori* survival and host interaction. Current evidence indicates that *H. pylori* infection is related to an increased likelihood of depleted iron storage (iron deficiency anemia) (Hudak, et al. 2017). Human iron metabolism is relevant to carcinogenesis (Torti, et al. 2018), and iron deficiency increases *H. pylori* virulence and the risk of GC (Hudak, et al. 2017). A large part of the *fecA-2* gene is deleted in hspAmerind strains. Mutations in *fecA-2* have been observed after *H. pylori* diversification in the Mongolian gerbil (Beckett, et al. 2018).

Correct disulfide bond (S-S) formation is critical in the folding of many secretory and membrane proteins in bacteria, including toxins, adherence factors, and components of secretory systems. The highly variable thiol:disulfide oxidoreductases of the Dsb (*d*i*s*ulfide *b*ond) family catalyze this step in the periplasm in gram-negative bacteria. DsbG/K of *H. pylori* (S6B Figure), a homolog of DsbG of *E. coli*, is secreted and affects the stomach epithelium (Kim, et al. 2002) and enables colonization (Kaakoush, et al. 2007). It can interact with and refold reduced HcpE (HP0235) (Lester, et al. 2015). DsbG/K acts on HopQ and helps HopQ-CEACAM interaction for the delivery of CagA in which another discriminatory SNP was found. Therefore, the loci identified by our GWAS are not only multifaceted, but can interact with and affect each other.

The discriminatory SNP with the lowest *P*-value was located upstream of *kch* (*trkA)* (S7 Figure), encoding a K+ channel protein essential for *H. pylori* colonization of the murine stomach. In human epithelial cells, various K^+^ channels are expressed, allowing adaptation to different needs in different organs (Heitzmann and Warth 2008). In the human gastric mucosa, K^+^ channel function is a prerequisite for acid secretion by parietal cells. In epithelial cells of the small intestine, K^+^ channels provide the driving force for electrogenic transport across the plasma membrane, and they are involved in cell volume regulation. Similarly, *H. pylori* may express this K^+^ channel in different ways for different needs in the two organs.

In addition to known virulence factors, our GWAS revealed three hitherto unrecognized virulence factor candidates: CtbP, Isp, and TriH (Figure 5). A likely common mechanism is interference with specific host proteins, a strategy shown for CagA, although there still is an element of competition for the small molecule NAD. They resemble several tumor virus oncoproteins, such as E1A of adenovirus and Tax of HTLV, which take over, by protein-protein interaction, the human cell protein network for survival of the infected cells (King, et al. 2018). *H. pylori*, an oncogenic bacterium, may use the same strategy as tumor viruses, competing with human cells over small molecules.

A previous GWAS comparing *H. pylori* from patients diagnosed with NAG and GC with a focus on the hpEurope population (Berthenet, et al. 2018) revealed 32 GC-associated loci. These genes mostly belonged to the *cag* pathogenicity island (PAI) and encoded outer membrane proteins, such as *babA*. In our GWAS, focusing on the hspEAsia population, we found none of these previously reported GC-associated loci. This discrepancy is likely due to two major differences. First, the preceding study compared GC and gastritis, while we compared GC and DU. GC develops from gastritis, whereas GC and DU diverge in an early stage. Second, the hpEurope and hspEAsia populations are genetically different. In particular, hpEurope includes 50–60% *cag* PAI-positive strains, whereas hspEAsia strains are nearly all *cag* PAI-positive (Olbermann, et al. 2010). The organization of *babABC* loci/alleles as well as other outer membrane proteins is quite different in hpEurope and hspEAsia strains.

The set of discriminatory SNPs identified in this study will be potentially applicable to personalized risk stratification in clinical settings for early-stage discrimination between GC and DU and for the selection of appropriate treatments. A recent study developed a high-throughput multiple allele detection assay (Zhang, et al. 2016). Incorporation of the discriminatory SNPs into such a technique will assist clinicians in diagnosis and clinical decision making.

In conclusion, our study revealed multifaceted genetic features of *H. pylori* associated with the pathogenesis of GC as compared to DU, and demonstrated the effectiveness of GWAS followed by regression-based prediction in distinguishing these *H. pylori*-related diseases, although the individual effect of each discriminatory SNP was not significant despite using the largest-to-date, but still limited in sample size dataset. Our analysis provided insights into the pathogenesis of GC and provided a basis for identifying a set of biomarkers for application in clinical settings.

## Materials and Methods

### Bacterial isolation and genome sequencing and assembly

*H. pylori* strains were isolated in Vietnam (from patients indicated for upper endoscopy at Cho Ray Hospital, Ho-Chi Minh and 108 Military Hospital, Hanoi) and in Oita, Japan, using standard culture methods. Briefly, homogenized antral biopsy specimens were inoculated on *H. pylori*-selective plates (Nissui Pharmaceutical Co., Ltd., Tokyo, Japan) and incubated at 37 °C in a microaerophilic condition for 3–10 days. Purple colonies that appeared were subcultured in Brucella Broth (Becton, Dickinson and Company, Sparks, MD) supplemented with 7% horse blood. DNA was extracted using a DNeasy Blood & Tissue Kit (Qiagen Inc., Valencia, CA). DNA concentrations were measured using a Quantus^™^ Fluorometer (Promega). High-throughput genome sequencing was performed on a HiSeq 2500 (2 × 100 or 2 × 150 paired-end reads) or MiSeq (2 × 300 paired-end reads) sequencer (Illumina, San Diego, CA), following the manufacturer’s instructions. Trimmomatic v0.35 was used to remove adapter sequences and low-quality bases from the raw short-read data (Bolger, et al. 2014). Trimmed reads were *de-novo* assembled to produce contigs using the SPAdes (v3.12.0) genome assembler with the “-careful” option to reduce mismatches in the assembly (Bankevich, et al. 2012). The minimum contig length was set to 200 bp.

Glycerol stocks of *H. pylori* strains isolated in Japan (from patients in different geographical areas of Japan, including Fukui, Okinawa, and Oita) were propagated on trypticase soy agar supplemented with 5% sheep blood (BD Biosciences, San Jose, CA) at 37 °C under microaerobic (5% O_2_) conditions in a HERAcell 150i CO_2_ incubator (Thermo Fisher Scientific, Waltham MA). *H. pylori* colonies were pooled, transferred into a Petri dish containing 40 mL of Brucella Broth supplemented with 10% fetal bovine serum (Sigma-Aldrich, St. Louis, MO), and incubated under agitation for three days. After incubation, the cells were harvested in 50-mL tubes, and frozen. Genomic DNA was extracted from the frozen cell pellets using Qiagen Genomic-tip 100/G, RNase A, Proteinase K, and Genomic DNA Buffer Set (all from Qiagen, Hilden, Germany), essentially following the protocol described in the Qiagen Genomic DNA Handbook. Genomic DNA was resuspended in TE buffer and sheared for library construction using a Covaris g-TUBE device according to the manufacturer’s instructions. A SMRTbell library was prepared using a SMRTbell Template Prep Kit 1.0 (Pacific Biosciences, Menlo Park, CA). DNA fragments larger than 17 kbp were size-selected using the BluePippin system (Sage Science, Beverly, MA). For each *H. pylori* strain, one SMRT cell was run on the PacBio RS II System with P6/C4 or P6/C4v2 chemistry and 360-min movies (Pacific Biosciences). SMRT sequencing data were analyzed using SMRT Analysis v2.3.0 through the SMRT Portal. Reads were assembled using RS_HGAP_Assembly.2. After the removal of overlapping ends, the chromosomal contig was reshaped to start from the ori-sequence. Thereafter, it was re-sequenced with RS_Resequencing.1 to create consensus sequences.

Assembled contigs of the 240 hspEAsia strains are available at https://figshare.com/s/2174da1fa20ae71c71e0. Sequencing and assembly statistics as well as other metadata of the newly sequenced strains are presented in Supplementary file 1, together with those of 405 other strains registered in public databases and initially analyzed in this study. Data of the newly PacBio sequenced strains were deposited in DDBJ and mirrored in NCBI under BioProject accession number PRJDB5843. Data of newly HiSeq sequenced strains isolated in Vietnam were deposited in DDBJ and mirrored in NCBI under BioProject accession number PRJDB10671, and those isolated in Japan were also deposited under BioProject accession number PRJDB10720, PRJNA215152, PRJNA215153, PRJNA246665, and PRJNA246666 (Supplementary Table 2).

### Population assignment of each strain

We inferred the population structure of 614 global strains in total, using chromosome painting and fineSTRUCTURE, as previously described (Lawson, et al. 2012). Briefly, a contig from each genome was initially mapped to the genome of strain 26695 as a reference, using Snippy v4.0.7 (https://github.com/tseemann/snippy). The Snippy-core function was used to create genome-wide haplotype data for all strains. Subsequently, ChromoPainter (v0.04) inferred chunks of DNA donated from a donor to a recipient for each recipient haplotype, and summarized the results into a co-ancestry matrix. Using this co-ancestry matrix, fineSTRUCTURE (v0.02) then clustered individuals by a Bayesian Markov chain Monte Carlo (MCMC) approach with 100,000 iterations for both the burn-in and the MCMC chain after the burn-in.

### GWAS

All isolates assigned to hspEAsia based on the fineSTRUCTURE results and for which clinical information of interest (GC or DU) was available were used for GWAS. First, a maximum-likelihood phylogenetic tree based on core-genome SNPs was reconstructed using PhyML (Guindon, et al. 2010), and the distribution of GC and DU in the tree was visualized using Phandango (Hadfield, et al. 2017). The tree is shown as mid-point-rooted. Core-genome SNPs were extracted based on mapping of each genome against that of the East Asian-type (hspEASia) *H. pylori* strain F57, using Snippy v4.0.7. We used strain F57 as a reference because it was isolated from a GC patient in Japan and its genome sequence has been determined by whole-genome shotgun sequencing (Kawai, et al. 2011).

Next, we conducted a pairwise genome alignment between each genome and strain F57 using progressiveMauve (Darling, et al. 2010), which enables the construction of positional homology alignments even for genomes with variable gene content and rearrangement. Subsequently, we combined all alignments into a multiple genome alignment in which each position corresponded to that of the strain F57 reference genome. Next, we extracted SNPs with ≤ 10% missing frequency and >5% minor allele frequency. We conducted a SNP GWAS based on a previous study (Berthenet, et al. 2018), in which a linear mixed regression model with the *bugwas* package (Earle, et al. 2016b) was used to control for population structure based on an n × n relatedness matrix calculated from bi-allelic SNPs. We also conducted a SNP GWAS in which the algorithm-factored spectrally transformed linear mixed model (FaST-LMM) implemented in pyseer (Lees, et al. 2018) was used to control for population structure from the same set of bi-allelic SNPs. A Q-Q plot was created using the R statistical program to assess the number and magnitude of observed associations between SNPs and disease (GC and DU) as compared to the association statistics expected under the null hypothesis of no association.

We also conducted a GWAS focusing on the presence or absence of specific genes rather than SNPs, based on pan-genome analysis using the Roary pipeline (Page, et al. 2015), although we did not conduct further analysis based on Q-Q plot assessment (see Results). A gene presence or absence matrix was used as an input of the linear mixed regression model implemented in the *bugwas* package.

### Discrimination between GC and DU using a set of SNPs identified by GWAS

Top hit SNPs deviating from the null hypothesis in the Q-Q plot and positively associated with GC were used to calculate a simple discriminatory score for each strain (Berthenet, et al. 2018). For each SNP (*j* = 1,…, *P*), where *P* indicates the number of top hit SNPs, when it has a nucleotide positively associated with GC, a variable *I_j_* is set to 1 if the strain has it or –1 if it does not. The discriminatory score of an individual strain *x*_i_ was calculated by taking the summation over all SNPs:

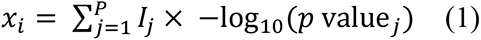

We next used a prediction approach as follows. We used a univariate logistic regression model to predict the probability of a strain *i* (*p_i_*) being isolated from a GC patient from the discriminatory score:

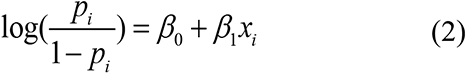

We then conducted a 2-fold cross-validation in which the logistic regression model was fit to a training dataset to estimate the regression coefficients, and the probability for each strain in a test dataset (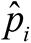) was predicted from *x*i. ROC curves were drawn from the true host disease status (GC or DU) and the predicted probability (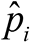) of each strain to calculate the AUC, determine the optimal cutoff value for the discrimination of GC and DU, and calculate the sensitivity and specificity of the discrimination, using the R package pROC (Robin, et al. 2011). Training and test datasets were prepared (see Results) to check the robustness of results.

### Analysis of amino acid and RNA changes at SNPs

Non-synonymous SNPs deviating from the null hypothesis were mapped on 3D structural models of their protein products (in strain F57 or 26695), using the automated homology modeling programs SWISS-MODEL (https://swissmodel.expasy.org/interactive) and PyMOL (Molecular Graphics System, v.1.2r3pre, Schrödinger, LLC). We also used KEGG (https://www.genome.jp/kegg/), UniProt (https://www.uniprot.org/), RCSB (https://www.rcsb.org/), SignalP-5.0 (http://www.cbs.dtu.dk/services/SignalP/), and cNLS Mapper (http://nls-mapper.iab.keio.ac.jp).

For intergenic SNPs deviating from the null hypothesis, we examined whether they were located in small regulatory RNAs previously identified in the reference strain 26695 (Sharma, et al. 2010) and registered in BSRD database (Li, et al. 2013) (http://kwanlab.bio.cuhk.edu.hk/BSRD/). M-fold (http://unafold.rna.albany.edu/?q=mfold) was used for secondary structure prediction.

## Acknowledgements

We thank Daniel Falush and Masao Ueki for discussion, and all the researchers worldwide that have whole-genome sequenced *H. pylori* isolates and made their data available to us. We are grateful to Karen Ottemann for comments on TlpC. Computational calculations were performed at the National Institute of Genetics (Japan). This work was supported by National Institute of Basic Biology (NIBB) Collaborative Research Program to I.K. This work was supported by Grants-in-Aid for Scientific Research from the Ministry of Education, Culture, Sports, Science and Technology (MEXT) (18KK0266 to Y.Y. and K.Y.; 25291080, 26113704, 17H04666 to I.K). This work was also supported by JSPS KAKENHI Grant Number 221S0002 and 16H06279 (PAGS).

## Competing interests

The authors declare that they have no competing interests.

---

**S1 Figure.**
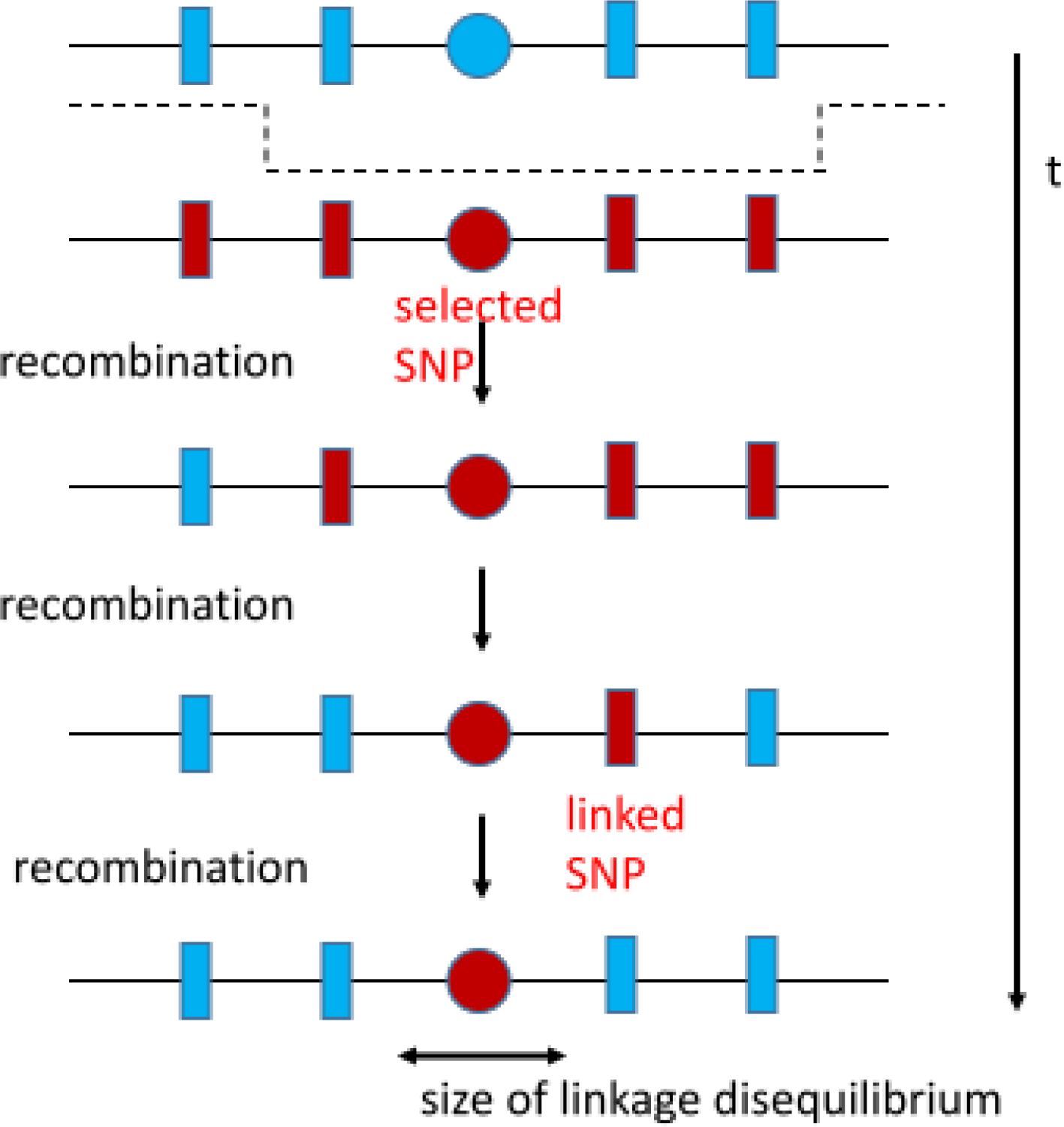
Schematic depiction of the decrease of linkage disequilibrium after repeated recombination events.

**S2 Figure.**
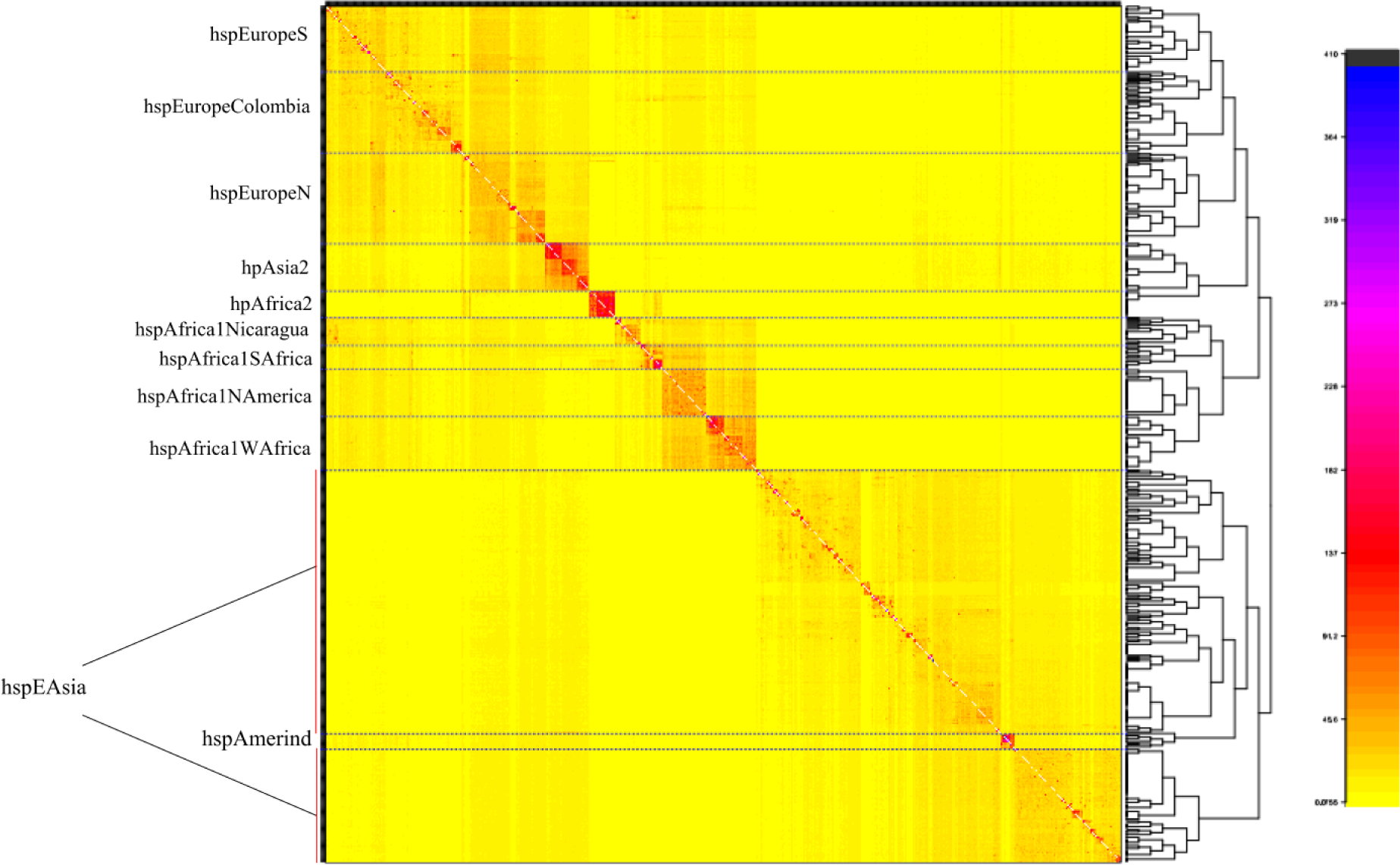
Population structure of 614 global *H. pylori* strains. The color of each cell of the co-ancestry matrix indicates the expected number of DNA chunks imported from a donor genome (column) to a recipient genome (row). Dashed lines separate the different populations. Strains belonging to hspEAsia are indicated by the red vertical lines. Detailed information on the *H. pylori* strains used in this analysis is shown in Table S1.

**S3 Figure.**
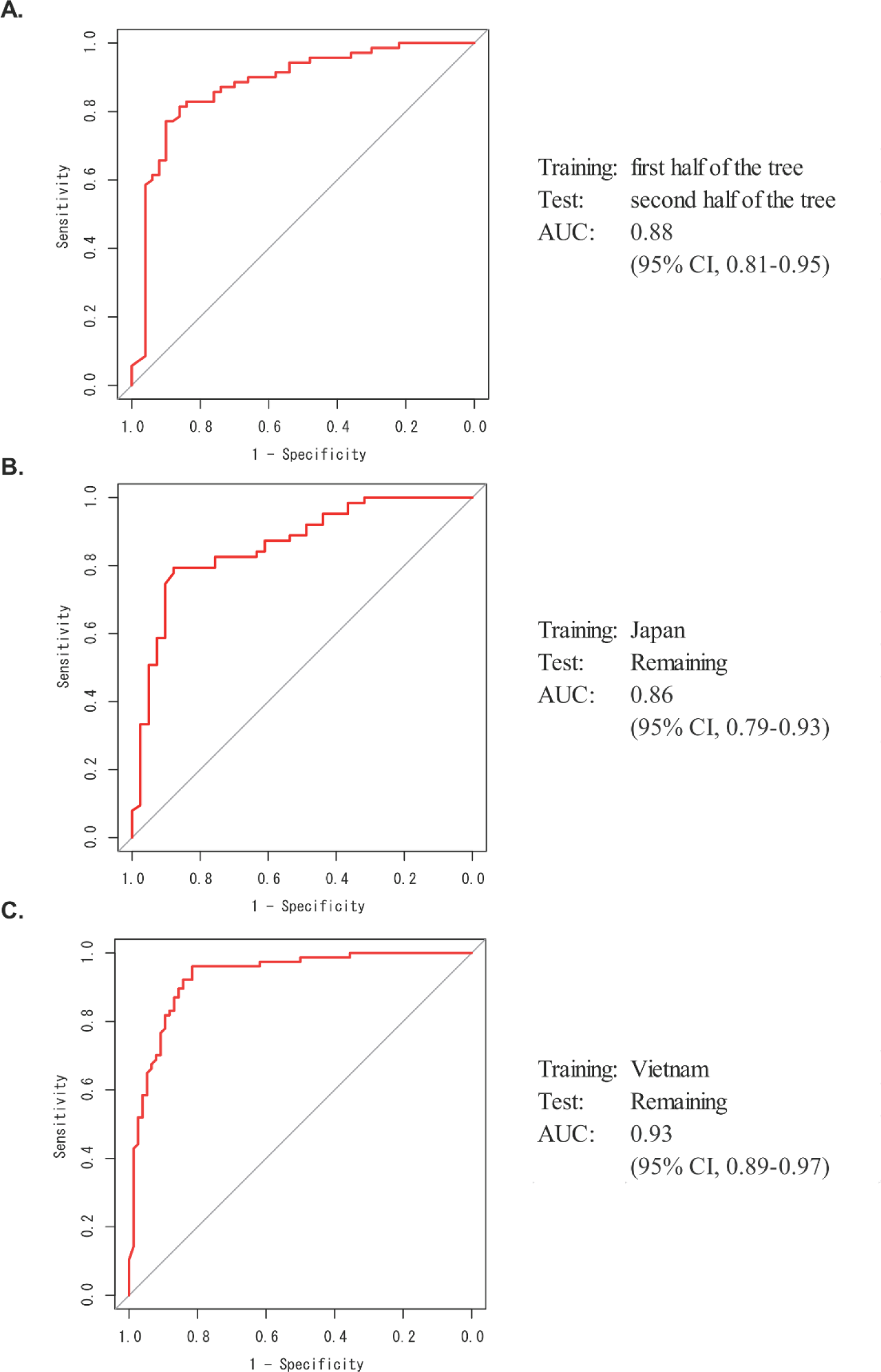
ROC curves representing performance of the discrimination. Sensitivity (y-axis) and 1 – specificity (x-axis) were calculated using three means of 2-fold cross validations and cutoffs of the predicted probability of being isolated from a GC patient for an strain.

**S4 Figure.**
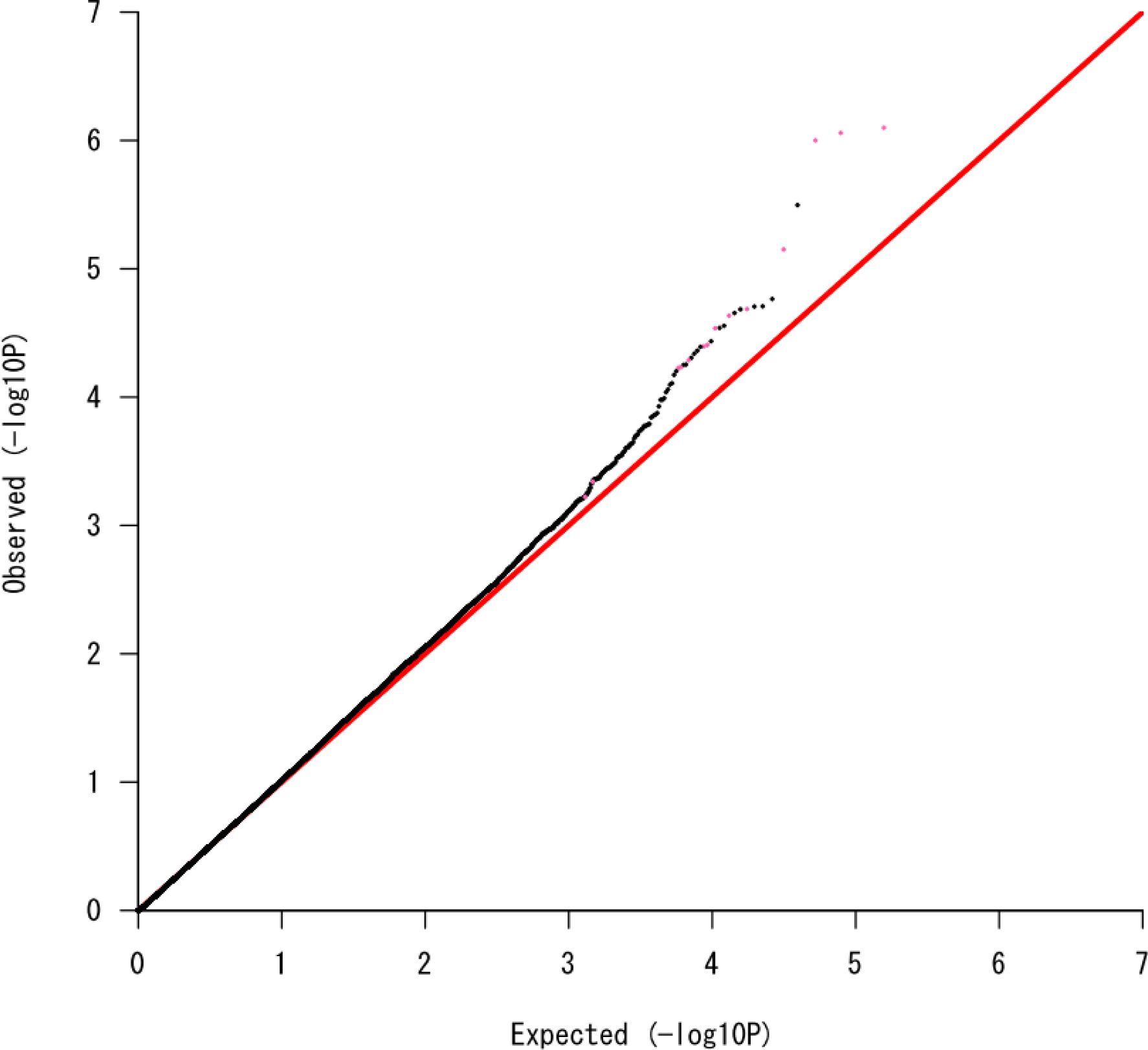
Q-Q plot to assess GWAS results using FaST-LMM. The 14 SNPs identified in Figure 2 are indicated in pink.

**S5 Figure.**
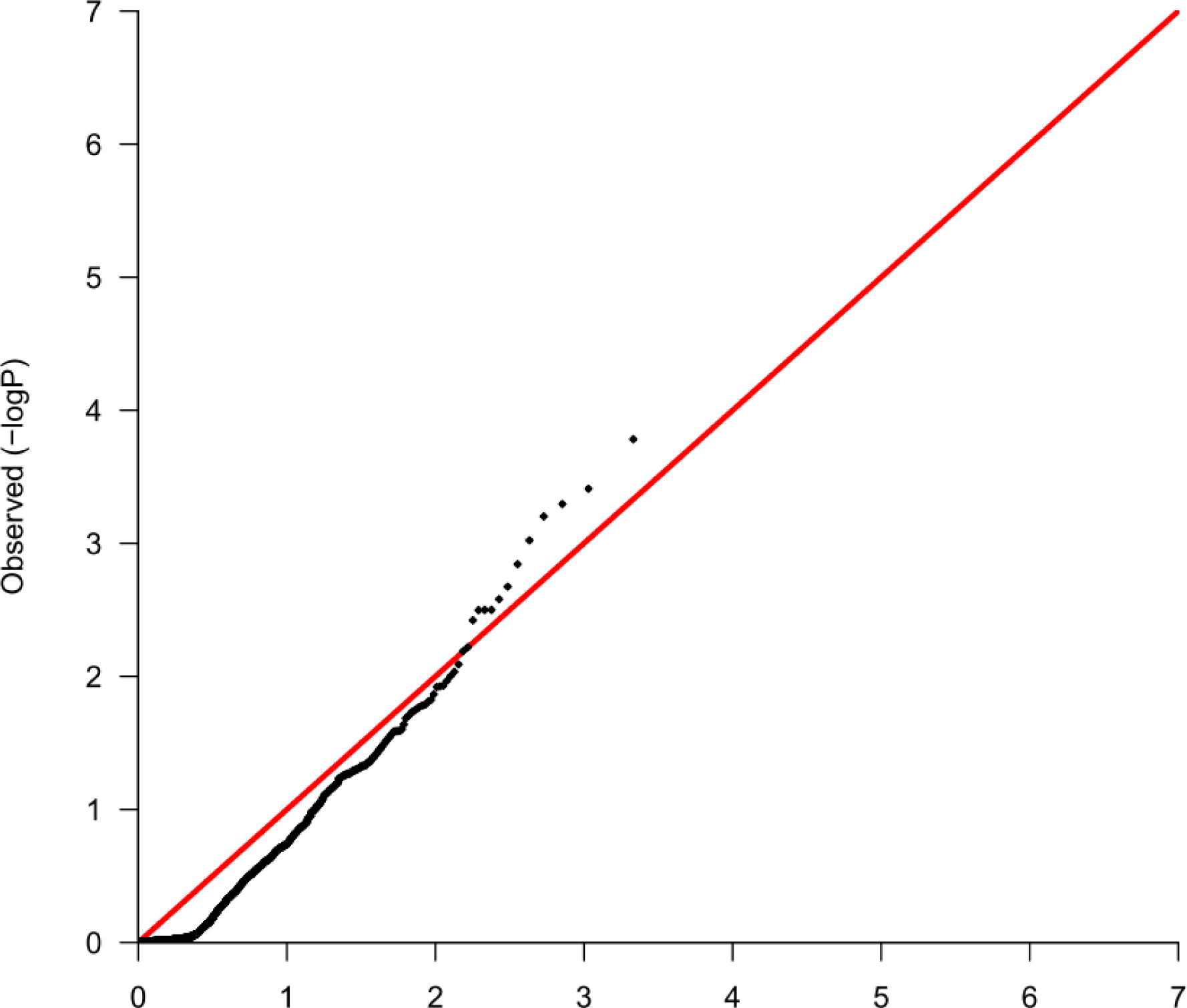
Q-Q plot to assess GWAS results regarding the presence or absence of a specific gene.

**S6 Figure.**
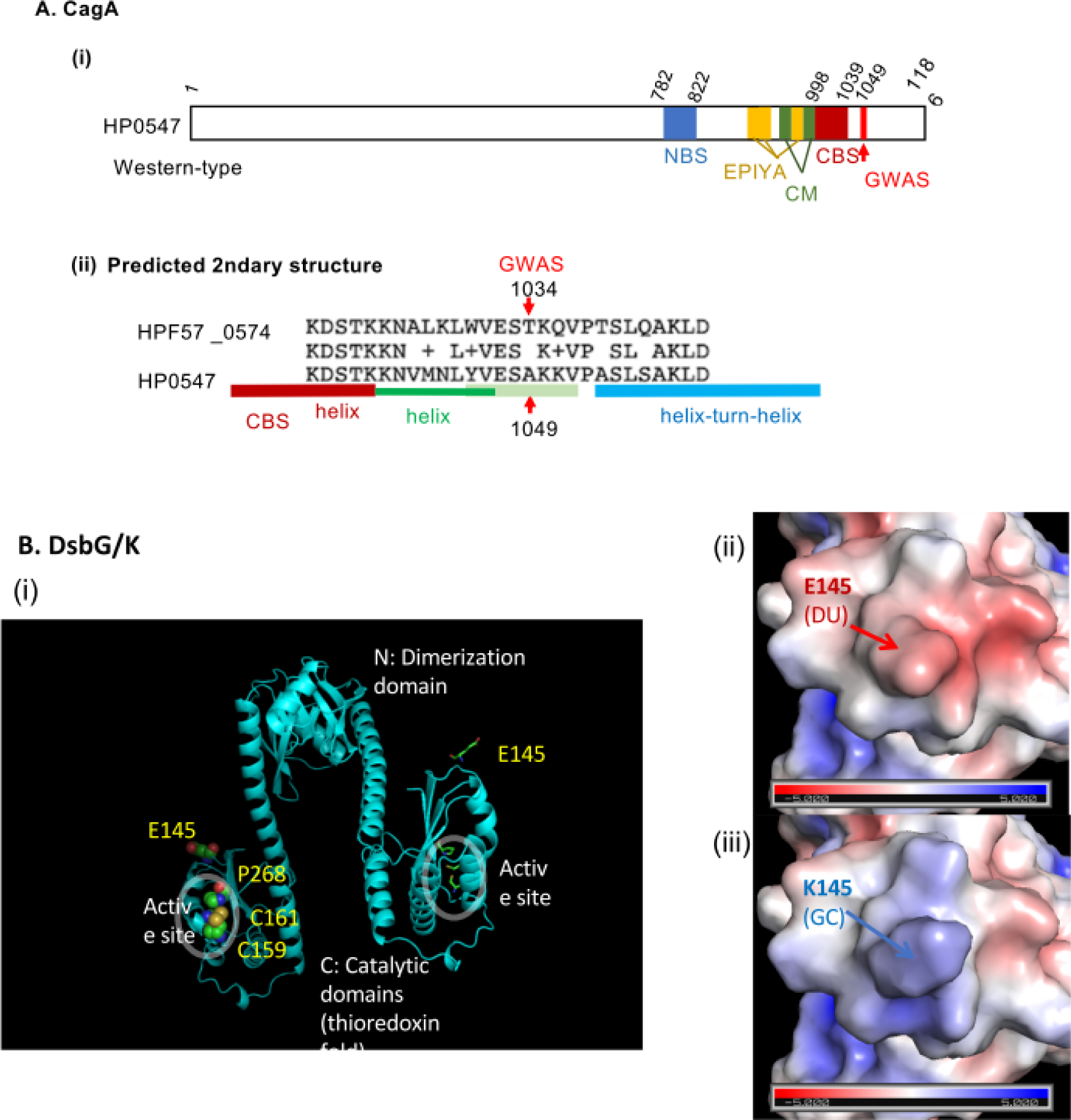
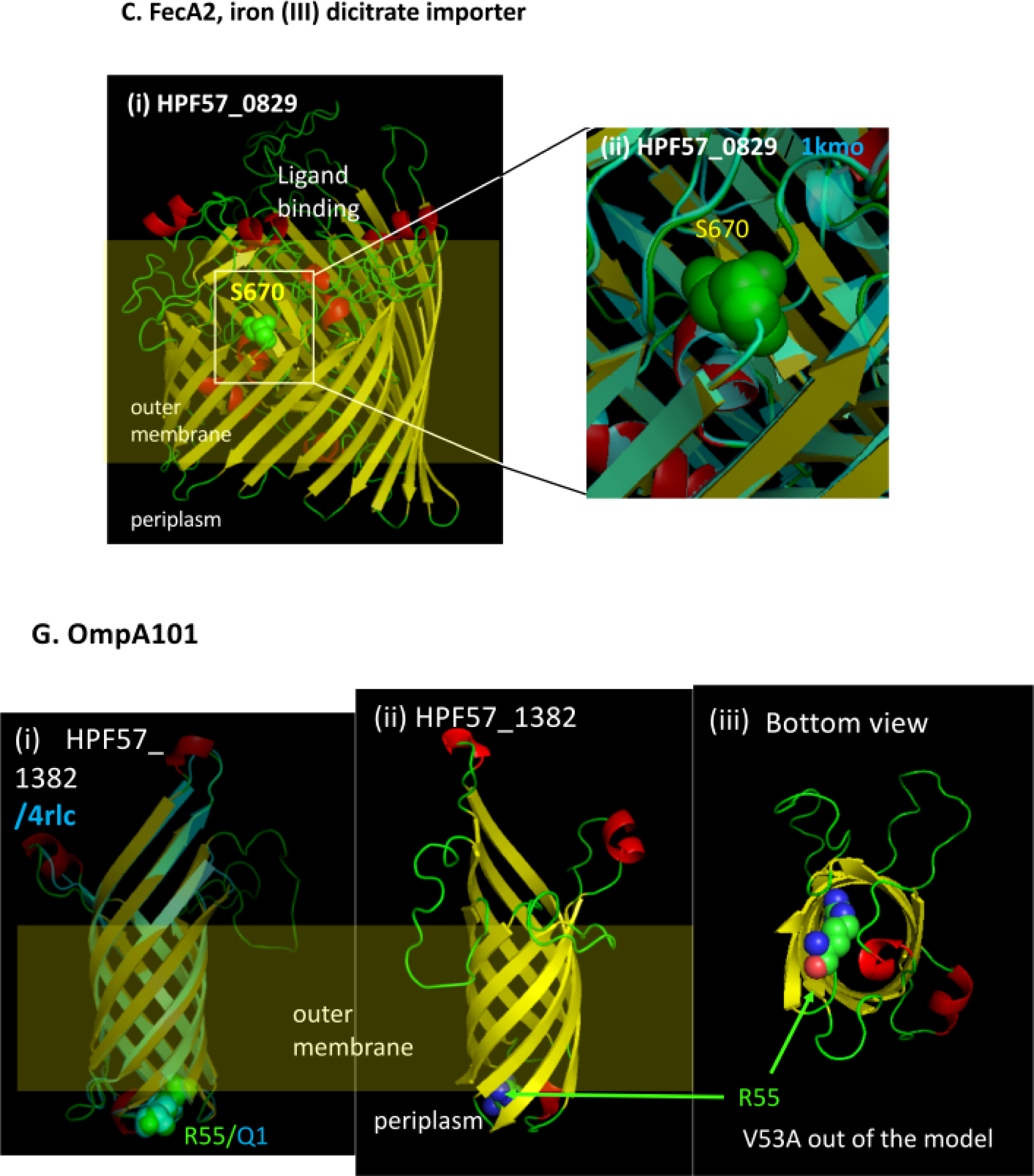
Proteins with amino-acid changes at a discriminatory SNP. **A.** CagA. **(i)** CagA (HPF57_0574, HP0547) has been extensively studied as an oncoprotein^1^. East-Asian-type CagA, including HPF57 _0574, likely evolved from Western-type, including HP0547, via Amerind-type as an intermediate ^2^. **(ii)** GWAS residue (A1034T/X/S) lies in L1029WVEST1034KQVPTS in F57_0574 and corresponds to A1049 in LVESA1049KKVPAS in HP0547. It is out of the solved structure (1–876 in HP0547). It is 12 aa away from the C-terminal binding sequence (CBS, 983–1023 in F57_0574, 998-1038 in HP0547), which is homologous to the solved N-terminal binding sequence (NBS)^3^ and is predicted to form a helix. This helix is predicted to extend through L1029 and, based on similarity with Lpg2149 (PDB 5dpo, 7bxf), further through EST1034KQV1037 until interruption by P1038. A pair of this helix (LKDSTKKNALKLWVEST1034KQV) can form a coiled coil. After P1038, a helix-turn-helix is predicted based on similarity with Arc repressor (PDB 1arr, 1arq, 1par). In summary, the GWAS site is likely to be between two helices and might itself lie on a helix. **B.** DsbG/K. **(i)** HPF57_0250 was modeled on HP0231 with 99% amino-acid identity (3tdg.1.A in PDB). The C-terminal domain (residues 132–265) has the thioredoxin-like fold with the catalytic CXXC motif and cis-Pro loop. The GWAS SNP site E145 (DU) lies at one side of the C-terminal domain distant from the CXXC motif. **(ii, iii)** Its mutation to K145 drastically changes the surface electric charge distribution and likely affects its function as a redox enzyme and its interaction with other components. **C.** FecA-2, a TonB-dependent outer membrane iron (III) dicitrate transporter. HPF57_0829 was modeled on 1kmo of PDB (FecA of *Escherichia coli*). S670 is predicted to be in a loop right above the wall of the barrel. The same location was found with models on 4aip and 4aiq (FrpB of *Neisseria meningitides*). **D.** OmpA101. HPF57_1382 was well modeled on *E. coli* OmpA structures and PDB 4rlc (N-terminal domain of *Pseudomonas aeruginosa* OprF) ((i)–(iii)). They are assembled into an 8-stranded β-barrel by the Bam machinery and may function in the uptake of small molecules^4^. V53A is on the second amino-acid residue from R55 at the N-terminus of the model, which lies in a loop at the bottom of the barrel. It is located after a predicted signal peptide (1–22) (SignalP-5.0). Based on the length from the barrel model, V53A corresponds to Q20 of *E. coli* OmpA, which is at the end of its signal sequence. V53A might affect ligand exit or own translocation through inner membrane.

1. Hatakeyama, M. Structure and function of Helicobacter pylori CagA, the first-identified bacterial protein involved in human cancer. *Proc Jpn Acad Ser B Phys Biol Sci* **93**, 196-219 (2017).
2. Furuta, Y., Yahara, K., Hatakeyama, M. & Kobayashi, I. Evolution of cagA oncogene of Helicobacter pylori through recombination. *PLoS One* **6**, e23499 (2011).
3. Hayashi, T. *et al*. Tertiary structure-function analysis reveals the pathogenic signaling potentiation mechanism of Helicobacter pylori oncogenic effector CagA. *Cell Host Microbe* **12**, 20-33 (2012).
4. Reusch, R.N. Insights into the structure and assembly of Escherichia coli outer membrane protein A. *FEBS J* **279**, 894-909 (2012).

**S7 Figure.**
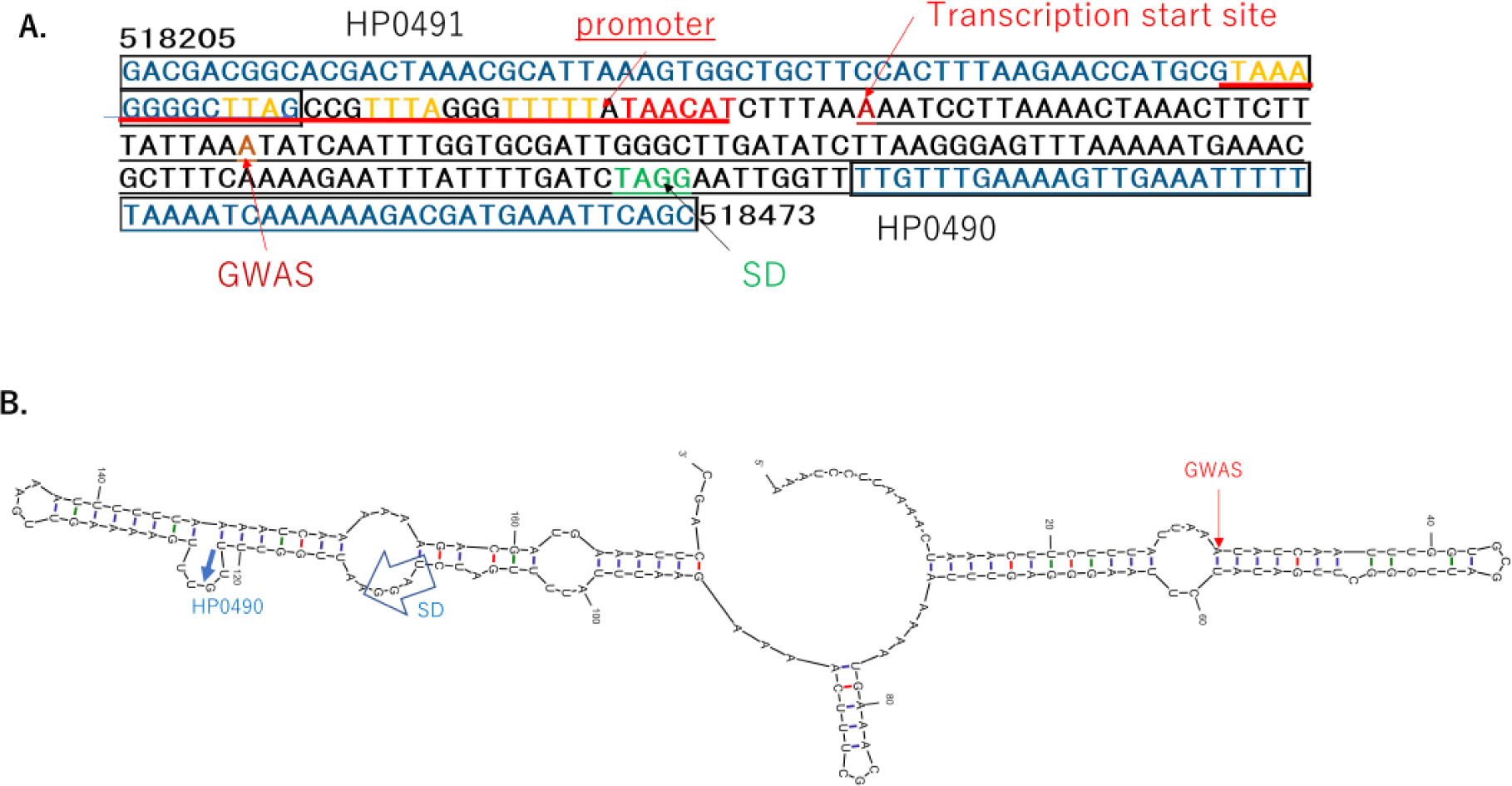
GWAS SNP upstream of *kch*, *<u>trkA</u>* (HP0490). **A.** Sequence. **B.** Predicted secondary structure of the 5′ part of the expected mRNA.

## References

1. Backert S, Bernegger S, Skorko-Glonek J, Wessler S. 2018. Extracellular HtrA serine proteases: An emerging new strategy in bacterial pathogenesis. Cell Microbiol 20:e12845.

2. Bankevich A, Nurk S, Antipov D, Gurevich AA, Dvorkin M, Kulikov AS, Lesin VM, Nikolenko SI, Pham S, Prjibelski AD, et al. 2012. SPAdes: a new genome assembly algorithm and its applications to single-cell sequencing. J Comput Biol 19:455–477.

3. Beckett AC, Loh JT, Chopra A, Leary S, Lin AS, McDonnell WJ, Dixon B, Noto JM, Israel DA, Peek RM, Jr., et al. 2018. Helicobacter pylori genetic diversification in the Mongolian gerbil model. PeerJ 6:e4803.

4. Bellesis AG, Jecrois AM, Hayes JA, Schiffer CA, Royer WE, Jr. 2018. Assembly of human C-terminal binding protein (CtBP) into tetramers. J Biol Chem 293:9101–9112.

5. Berthenet E, Yahara K, Thorell K, Pascoe B, Meric G, Mikhail JM, Engstrand L, Enroth H, Burette A, Megraud F, et al. 2018. A GWAS on Helicobacter pylori strains points to genetic variants associated with gastric cancer risk. BMC Biol 16:84.

6. Bischler T, Tan HS, Nieselt K, Sharma CM. 2015. Differential RNA-seq (dRNA-seq) for annotation of transcriptional start sites and small RNAs in Helicobacter pylori. Methods 86:89–101.

7. Blair KM, Mears KS, Taylor JA, Fero J, Jones LA, Gafken PR, Whitney JC, Salama NR. 2018. The Helicobacter pyloricell shape promoting protein Csd5 interacts with the cell wall, MurF, and the bacterial cytoskeleton. Mol Microbiol 110:114–127.

8. Bolger AM, Lohse M, Usadel B. 2014. Trimmomatic: a flexible trimmer for Illumina sequence data. Bioinformatics 30:2114–2120.

9. Breurec S, Guillard B, Hem S, Brisse S, Dieye FB, Huerre M, Oung C, Raymond J, Tan TS, Thiberge JM, et al. 2011. Evolutionary history of Helicobacter pylori sequences reflect past human migrations in Southeast Asia. PLoS One 6:e22058.

10. Cheng H, Zhang N, Pati D. 2020. Cohesin subunit RAD21: From biology to disease. Gene 758:144966.

11. Chewapreecha C, Marttinen P, Croucher NJ, Salter SJ, Harris SR, Mather AE, Hanage WP, Goldblatt D, Nosten FH, Turner C, et al. 2014. Comprehensive identification of single nucleotide polymorphisms associated with beta-lactam resistance within pneumococcal mosaic genes. PLoS Genet 10:e1004547.

12. Chinnadurai G. 2009. The transcriptional corepressor CtBP: a foe of multiple tumor suppressors. Cancer Res 69:731–734.

13. Correa P, Haenszel W, Cuello C, Tannenbaum S, Archer M. 1975. A model for gastric cancer epidemiology. Lancet 2:58–60.

14. Darling AE, Mau B, Perna NT. 2010. progressiveMauve: multiple genome alignment with gene gain, loss and rearrangement. PLoS One 5:e11147.

15. Earle SG, Wu C-H, Charlesworth J, Stoesser N, Gordon NC, Walker TM, Spencer CCA, Iqbal Z, Clifton DA, Hopkins KL, et al. 2016a. Identifying lineage effects when controlling for population structure improves power in bacterial association studies. Nature Microbiology 1:Article number: 16041

16. Earle SG, Wu CH, Charlesworth J, Stoesser N, Gordon NC, Walker TM, Spencer CCA, Iqbal Z, Clifton DA, Hopkins KL, et al. 2016b. Identifying lineage effects when controlling for population structure improves power in bacterial association studies. Nat Microbiol 1:16041.

17. El Khadir M, Alaoui Boukhris S, Benajah DA, El Rhazi K, Ibrahimi SA, El Abkari M, Harmouch T, Nejjari C, Mahmoud M, Benlemlih M, et al. 2017. VacA and CagA Status as Biomarker of Two Opposite End Outcomes of Helicobacter pylori Infection (Gastric Cancer and Duodenal Ulcer) in a Moroccan Population. PLoS One 12:e0170616.

18. Falush D. 2016. Bacterial genomics: Microbial GWAS coming of age. Nat Microbiol 1:16059.

19. Falush D, Bowden R. 2006. Genome-wide association mapping in bacteria? Trends Microbiol 14:353–355.

20. Furuta Y, Yahara K, Hatakeyama M, Kobayashi I. 2011. Evolution of cagA oncogene of Helicobacter pylori through recombination. PLoS One 6:e23499.

21. Guindon S, Dufayard JF, Lefort V, Anisimova M, Hordijk W, Gascuel O. 2010. New algorithms and methods to estimate maximum-likelihood phylogenies: assessing the performance of PhyML 3.0. Syst Biol 59:307–321.

22. Hadfield J, Croucher NJ, Goater RJ, Abudahab K, Aanensen DM, Harris SR. 2017. Phandango: an interactive viewer for bacterial population genomics. bioRxiv.

23. Hansson LE, Nyren O, Hsing AW, Bergstrom R, Josefsson S, Chow WH, Fraumeni JF, Jr., Adami HO. 1996. The risk of stomach cancer in patients with gastric or duodenal ulcer disease. N Engl J Med 335:242–249.

24. Hatakeyama M. 2017. Structure and function of Helicobacter pylori CagA, the first-identified bacterial protein involved in human cancer. Proc Jpn Acad Ser B Phys Biol Sci 93:196–219.

25. Hauryliuk V, Atkinson GC, Murakami KS, Tenson T, Gerdes K. 2015. Recent functional insights into the role of (p)ppGpp in bacterial physiology. Nat Rev Microbiol 13:298–309.

26. Heitzmann D, Warth R. 2008. Physiology and pathophysiology of potassium channels in gastrointestinal epithelia. Physiol Rev 88:1119–1182.

27. Hudak L, Jaraisy A, Haj S, Muhsen K. 2017. An updated systematic review and meta-analysis on the association between Helicobacter pylori infection and iron deficiency anemia. Helicobacter 22.

28. Jaillard M, Lima L, Tournoud M, Mahe P, van Belkum A, Lacroix V, Jacob L. 2018. A fast and agnostic method for bacterial genome-wide association studies: Bridging the gap between k-mers and genetic events. PLoS Genet 14:e1007758.

29. Jetten AM. 2019. Emerging Roles of GLI-Similar Kruppel-like Zinc Finger Transcription Factors in Leukemia and Other Cancers. Trends Cancer 5:547–557.

30. Kaakoush NO, Kovach Z, Mendz GL. 2007. Potential role of thiol:disulfide oxidoreductases in the pathogenesis of Helicobacter pylori. FEMS Immunol Med Microbiol 50:177–183.

31. Kawai M, Furuta Y, Yahara K, Tsuru T, Oshima K, Handa N, Takahashi N, Yoshida M, Azuma T, Hattori M, et al. 2011. Evolution in an oncogenic bacterial species with extreme genome plasticity: Helicobacter pylori East Asian genomes. BMC Microbiol 11:104.

32. Keilberg D, Steele N, Fan S, Yang C, Zavros Y, Ottemann KM. 2020. Gastric metabolomics detects H. pylori correlated loss of numerous metabolites in both the corpus and antrum. Infect Immun.

33. Kelleher JE, Daniel AS, Murray NE. 1991. Mutations that confer de novo activity upon a maintenance methyltransferase. J Mol Biol 221:431–440.

34. Kennaway CK, Obarska-Kosinska A, White JH, Tuszynska I, Cooper LP, Bujnicki JM, Trinick J, Dryden DT. 2009. The structure of M.EcoKI Type I DNA methyltransferase with a DNA mimic antirestriction protein. Nucleic Acids Res 37:762–770.

35. Kim N, Weeks DL, Shin JM, Scott DR, Young MK, Sachs G. 2002. Proteins released by Helicobacter pylori in vitro. J Bacteriol 184:6155–6162.

36. King CR, Zhang A, Tessier TM, Gameiro SF, Mymryk JS. 2018. Hacking the Cell: Network Intrusion and Exploitation by Adenovirus E1A. mBio 9.

37. Kusters JG, van Vliet AH, Kuipers EJ. 2006. Pathogenesis of Helicobacter pylori infection. Clin Microbiol Rev 19:449–490.

38. Lawson DJ, Hellenthal G, Myers S, Falush D. 2012. Inference of population structure using dense haplotype data. PLoS Genet 8:e1002453.

39. Lee JH, Jun SH, Baik SC, Kim DR, Park JY, Lee YS, Choi CH, Lee JC. 2012. Prediction and screening of nuclear targeting proteins with nuclear localization signals in Helicobacter pylori. J Microbiol Methods 91:490–496.

40. Lees JA, Galardini M, Bentley SD, Weiser JN, Corander J. 2018. pyseer: a comprehensive tool for microbial pangenome-wide association studies. Bioinformatics 34:4310–4312.

41. Lester J, Kichler S, Oickle B, Fairweather S, Oberc A, Chahal J, Ratnayake D, Creuzenet C. 2015. Characterization of Helicobacter pylori HP0231 (DsbK): role in disulfide bond formation, redox homeostasis and production of Helicobacter cystein-rich protein HcpE. Mol Microbiol 96:110–133.

42. Li L, Huang D, Cheung MK, Nong W, Huang Q, Kwan HS. 2013. BSRD: a repository for bacterial small regulatory RNA. Nucleic Acids Res 41:D233–238.

43. Li X, Wang J, Shi Y. 2011. Structural mechanisms of DIAP1 auto-inhibition and DIAP1-mediated inhibition of drICE. Nat Commun 2:408.

44. Linz B, Balloux F, Moodley Y, Manica A, Liu H, Roumagnac P, Falush D, Stamer C, Prugnolle F, van der Merwe SW, et al. 2007. An African origin for the intimate association between humans and Helicobacter pylori. Nature 445:915–918.

45. Liu YP, Tang Q, Zhang JZ, Tian LF, Gao P, Yan XX. 2017. Structural basis underlying complex assembly and conformational transition of the type I R-M system. Proc Natl Acad Sci U S A 114:11151–11156.

46. Lu H, Hsu PI, Graham DY, Yamaoka Y. 2005. Duodenal ulcer promoting gene of Helicobacter pylori. Gastroenterology 128:833–848.

47. Machuca MA, Johnson KS, Liu YC, Steer DL, Ottemann KM, Roujeinikova A. 2017. Helicobacter pylori chemoreceptor TlpC mediates chemotaxis to lactate. Sci Rep 7:14089.

48. Mechold U, Potrykus K, Murphy H, Murakami KS, Cashel M. 2013. Differential regulation by ppGpp versus pppGpp in Escherichia coli. Nucleic Acids Res 41:6175–6189.

49. Nishimori I, Minakuchi T, Kohsaki T, Onishi S, Takeuchi H, Vullo D, Scozzafava A, Supuran CT. 2007. Carbonic anhydrase inhibitors: the beta-carbonic anhydrase from Helicobacter pylori is a new target for sulfonamide and sulfamate inhibitors. Bioorg Med Chem Lett 17:3585–3594.

50. Olbermann P, Josenhans C, Moodley Y, Uhr M, Stamer C, Vauterin M, Suerbaum S, Achtman M, Linz B. 2010. A global overview of the genetic and functional diversity in the Helicobacter pylori cag pathogenicity island. PLoS Genet 6:e1001069.

51. Page AJ, Cummins CA, Hunt M, Wong VK, Reuter S, Holden MT, Fookes M, Falush D, Keane JA, Parkhill J. 2015. Roary: rapid large-scale prokaryote pan genome analysis. Bioinformatics 31:3691–3693.

52. Peek RM, Jr., Blaser MJ. 1997. Pathophysiology of Helicobacter pylori-induced gastritis and peptic ulcer disease. Am J Med 102:200–207.

53. Robin X, Turck N, Hainard A, Tiberti N, Lisacek F, Sanchez JC, Muller M. 2011. pROC: an open-source package for R and S+ to analyze and compare ROC curves. BMC Bioinformatics 12:77.

54. Rodionov DA, Arzamasov AA, Khoroshkin MS, Iablokov SN, Leyn SA, Peterson SN, Novichkov PS, Osterman AL. 2019. Micronutrient Requirements and Sharing Capabilities of the Human Gut Microbiome. Front Microbiol 10:1316.

55. San JE, Baichoo S, Kanzi A, Moosa Y, Lessells R, Fonseca V, Mogaka J, Power R, de Oliveira T. 2019. Current Affairs of Microbial Genome-Wide Association Studies: Approaches, Bottlenecks and Analytical Pitfalls. Front Microbiol 10:3119.

56. Sharma CM, Hoffmann S, Darfeuille F, Reignier J, Findeiss S, Sittka A, Chabas S, Reiche K, Hackermuller J, Reinhardt R, et al. 2010. The primary transcriptome of the major human pathogen Helicobacter pylori. Nature 464:250–255.

57. Sheppard SK, Didelot X, Meric G, Torralbo A, Jolley KA, Kelly DJ, Bentley SD, Maiden MC, Parkhill J, Falush D. 2013. Genome-wide association study identifies vitamin B5 biosynthesis as a host specificity factor in Campylobacter. Proc Natl Acad Sci U S A 110:11923–11927.

58. Shiota S, Matsunari O, Watada M, Hanada K, Yamaoka Y. 2010. Systematic review and meta-analysis: the relationship between the Helicobacter pylori dupA gene and clinical outcomes. Gut Pathog 2:13.

59. Sohn J, Grant RA, Sauer RT. 2010. Allostery is an intrinsic property of the protease domain of DegS: implications for enzyme function and evolution. J Biol Chem 285:34039–34047.

60. Stingl K, Brandt S, Uhlemann EM, Schmid R, Altendorf K, Zeilinger C, Ecobichon C, Labigne A, Bakker EP, de Reuse H. 2007. Channel-mediated potassium uptake in Helicobacter pylori is essential for gastric colonization. EMBO J 26:232–241.

61. Supuran CT, Capasso C. 2016. New light on bacterial carbonic anhydrases phylogeny based on the analysis of signal peptide sequences. J Enzyme Inhib Med Chem 31:1254–1260.

62. Suzuki M, Shibayama K, Yahara K. 2016. A genome-wide association study identifies a horizontally transferred bacterial surface adhesin gene associated with antimicrobial resistant strains. Sci Rep 6:37811.

63. Takahashi-Kanemitsu A, Knight CT, Hatakeyama M. 2020. Molecular anatomy and pathogenic actions of Helicobacter pylori CagA that underpin gastric carcinogenesis. Cell Mol Immunol 17:50–63.

64. Thorell K, Yahara K, Berthenet E, Lawson DJ, Mikhail J, Kato I, Mendez A, Rizzato C, Bravo MM, Suzuki R, et al. 2017. Rapid evolution of distinct Helicobacter pylori subpopulations in the Americas. PLoS Genet 13:e1006546.

65. Torti SV, Manz DH, Paul BT, Blanchette-Farra N, Torti FM. 2018. Iron and Cancer. Annu Rev Nutr 38:97–125.

66. Tsuruta O, Yokoyama H, Fujii S. 2012. A new crystal lattice structure of Helicobacter pylori neutrophil-activating protein (HP-NAP). Acta Crystallogr Sect F Struct Biol Cryst Commun 68:134–140.

67. Uemura N, Okamoto S, Yamamoto S, Matsumura N, Yamaguchi S, Yamakido M, Taniyama K, Sasaki N, Schlemper RJ. 2001. Helicobacter pylori infection and the development of gastric cancer. N Engl J Med 345:784–789.

68. Waldman T. 2020. Emerging themes in cohesin cancer biology. Nat Rev Cancer 20:504–515.

69. Wilken C, Kitzing K, Kurzbauer R, Ehrmann M, Clausen T. 2004. Crystal structure of the DegS stress sensor: How a PDZ domain recognizes misfolded protein and activates a protease. Cell 117:483–494.

70. Winkler W, Nahvi A, Breaker RR. 2002. Thiamine derivatives bind messenger RNAs directly to regulate bacterial gene expression. Nature 419:952–956.

71. Yahara K, Furuta Y, Morimoto S, Kikutake C, Komukai S, Matelska D, Dunin-Horkawicz S, Bujnicki JM, Uchiyama I, Kobayashi I. 2016. Genome-wide survey of codons under diversifying selection in a highly recombining bacterial species, Helicobacter pylori. DNA Res 23:135–143.

72. Yahara K, Furuta Y, Oshima K, Yoshida M, Azuma T, Hattori M, Uchiyama I, Kobayashi I. 2013. Chromosome painting in silico in a bacterial species reveals fine population structure. Mol Biol Evol 30:1454–1464.

73. Yahara K, Meric G, Taylor AJ, de Vries SP, Murray S, Pascoe B, Mageiros L, Torralbo A, Vidal A, Ridley A, et al. 2017. Genome-wide association of functional traits linked with Campylobacter jejuni survival from farm to fork. Environ Microbiol 19:361–380.

74. Yamamoto K, Yamanaka Y, Shimada T, Sarkar P, Yoshida M, Bhardwaj N, Watanabe H, Taira Y, Chatterji D, Ishihama A. 2018. Altered Distribution of RNA Polymerase Lacking the Omega Subunit within the Prophages along the Escherichia coli K-12 Genome. mSystems 3.

75. Yamaoka Y. 2010. Mechanisms of disease: Helicobacter pylori virulence factors. Nat Rev Gastroenterol Hepatol 7:629–641.

76. Yano H, Alam MZ, Rimbara E, Shibata TF, Fukuyo M, Furuta Y, Nishiyama T, Shigenobu S, Hasebe M, Toyoda A, et al. 2020. Networking and Specificity-Changing DNA Methyltransferases in Helicobacter pylori. Front Microbiol 11:1628.

77. Zanotti G, Papinutto E, Dundon W, Battistutta R, Seveso M, Giudice G, Rappuoli R, Montecucco C. 2002. Structure of the neutrophil-activating protein from Helicobacter pylori. J Mol Biol 323:125–130.

78. Zhang Y, Zhao F, Kong M, Wang S, Nan L, Hu B, Olszewski MA, Miao Y, Ji D, Jiang W, et al. 2016. Validation of a High-Throughput Multiplex Genetic Detection System for Helicobacter pylori Identification, Quantification, Virulence, and Resistance Analysis. Front Microbiol 7:1401.

79. Zurawa-Janicka D, Wenta T, Jarzab M, Skorko-Glonek J, Glaza P, Gieldon A, Ciarkowski J, Lipinska B. 2017. Structural insights into the activation mechanisms of human HtrA serine proteases. Arch Biochem Biophys 621:6–23.

